# Low-cost camera-based estrous tracking enables transgenesis in *Peromyscus leucopus*, the primary reservoir for Lyme disease

**DOI:** 10.1101/2023.10.20.563285

**Authors:** Joanna Buchthal, Emma J. Chory, Zachary Hill, Christy Dennison, Boqiang Tu, Rick P. Wierenga, Çağrı Çevrim, Stefan Golas, Sam R. Telford, Kara L. McKinley, Rudolf Jaenisch, Styliani Markoulaki, Kevin M. Esvelt

**Author notes:** Designates equal contribution (Joanna Buchthal, Emma J. Chory).

## Abstract

CRISPR/Cas9 technology has revolutionized the production of animal models by reducing experimental timelines, slashing costs and streamlining gene editing, leading to a rapid expansion in the number of unique models for the study of human disease and gene function. However, most non-model animals, many of which are important in cancer and aging research, remain recalcitrant to genome engineering due to our limited understanding of their reproductive biology. Many wild rodents that transmit human diseases remain particularly challenging to engineer due to low pregnancy rates, the lack of external copulatory plugs, and susceptibility to premature termination of pregnancy. Here, we present low-cost activity-based estrous tracking for the efficient generation of timed pregnant and pseudopregnant white-footed mice and extend this tracking method to both lab mice and hamsters. Leveraging this technology, we demonstrate the generation of engineered *Peromyscus leucopus*, the primary reservoir for Lyme disease-causing bacteria and a putative model organism for studies of aging. These tools have broad implications for biomedical research and ecological engineering.

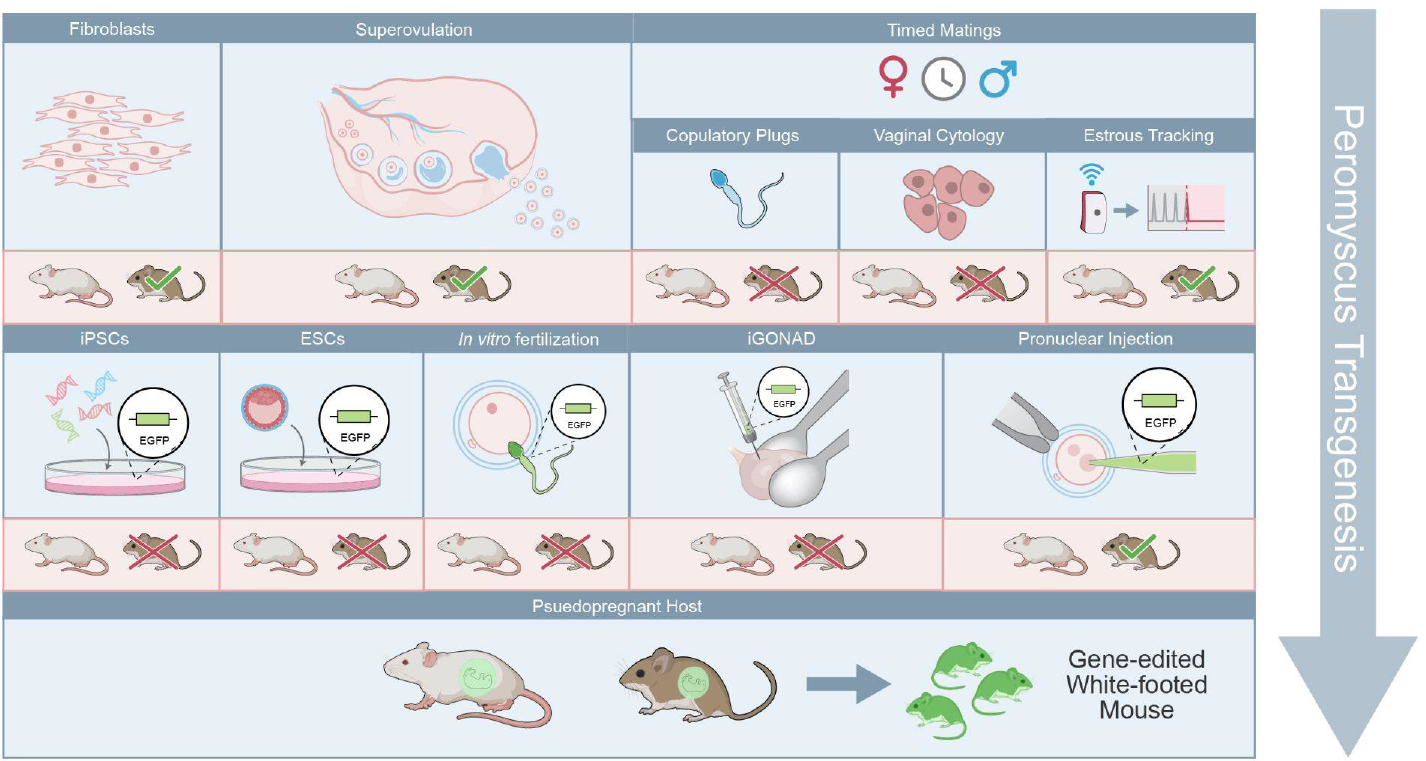

## INTRODUCTION

The deer mouse, also known as the white-footed or oldfield mouse, is the most abundant and diverse indigenous group of mammals in the United States. With a geographic range that spans from Alaska and northern Canada to western Panama, the *Peromyscus* genus includes a wealth of genetic diversity, prompting over a century of scientific observations. Consequently, more is known about the mammalian genus *Peromyscus* than any other group of small rodents besides the laboratory mouse and rat (Bedford & Hoekstra, 2015). Having contributed to the widespread understanding of evolution, ecology, reproductive biology, endocrinology and behavioral neuroscience, the deer mouse has been dubbed ‘the *Drosophila* of North American mammalogy (Dewey & Dawson, 2001).

The white-footed mouse *Peromyscus leucopus* is of particular interest due to its role in disease transmission, including Lyme disease, Babesiosis (Spielman et al., 1981), and Anaplasmosis (Telford et al., 1996), among others (Barbour, 2017). With much longer lifespans and patterns of genetic diversity more closely corresponding to humans than those of laboratory mice (Havighorst et al., 2017), *Peromyscus* have considerable potential for modeling human aging, disease, and pathology. Given their importance as an ecological model and their potential value to biomedical research, the ability to generate genetically engineered *Peromyscus* models would have widespread appeal. However, this genus has remained recalcitrant to engineering.

Since the first transgenic laboratory mice were engineered with viral DNA (Jaenisch & Mintz, 1974), scientists have developed increasingly powerful strategies for generating new animal models. Stem cell-based transgenesis technologies, which leverage mouse embryonic stem cells (mESCs), induced pluripotent stem cells (iPSCs)(Okita et al., 2007), and more recently, CRISPR/Cas9 for zygote engineering (Wu et al., 2013) constitute some of the most robust engineering approaches. Other techniques include i-GONAD, a method for delivering DNA via intraoviductal injection followed by *in vivo* electroporation (Ohtsuka et al., 2018), as well *ex vivo* and *in vivo* genome editing using recombinant adeno-associated viruses (rAAVs)(Yoon et al., 2018) and the lentiviral vector-mediated transduction of spermatozoa (Nagano et al., 2002) via in vitro fertilization. Some of these techniques, which were originally developed in *Mus musculus*, have been successfully applied to engineering new species (Fan et al., 2014; Hamra et al., 2002; Kobayashi et al., 2018). However, despite these advancements, these techniques have yet to be successfully applied to *Peromyscus*, and many other non-model rodents remain resistant to such engineering efforts.

Diverging from *Mus* over 25 million years ago, Peromyscus display a range of biological differences, from extended gestational timelines to physical differences in reproductive organs (Bedford & Hoekstra, 2015; Dewsbury et al., 1977; Dewsbury & Lanier, 1976; Hartung & Dewsbury, 1978). To be sure, these differences have confounded researchers who seek to develop reproductive protocols to engineer *Peromyscus*.

Attempts to apply conventional *Mus* reproductive protocols to *Peromyscus* have been markedly unsuccessful. Without visible copulatory plugs (Hartung & Dewsbury, 1978), researchers cannot use “plug checking” to verify mating, a commonly employed technique for mating detection in *Mus*, and the primary approach for confirming pseudopregnancy. As a result, researchers have turned to vaginal lavage for estrus and mating detection. However, by lavage, the *Peromsycus* estrous cycle appears highly variable, with estrous cycles lasting between 2 and 8+ days, despite having an assumed estrous cycle of 4 days (mode = 4, median = 4.9, average = 6)(Dewsbury et al., 1977). Further, in *Peromyscus*, the use of vaginal lavage to detect sperm appears to be minimally predictive of fertilization events, as researchers have routinely found sperm in vaginal smears from nongravid *P. leucopus* (Dewsbury & Lanier, 1976). Consequently, timed mating has been reported to be highly inefficient in *P. leucopus*, with a published pregnancy rate of 20% in females mated with males to satiety (Dewsbury & Lanier, 1976). In order to increase pregnancy rates (and thus embryo yields), other researchers have combined superovulation, timed mating and artificial insemination, but still only produced a fraction of the embryos that are typically harvested by superovulation alone in *Mus* (Veres et al., 2012). Finally, since mating events in *Peromyscus* regularly occur without giving rise to pregnancy, more recent advances such as mating detection with cameras are also unlikely to be successful in *Peromyscus* (Doroba & Sears, 2010; Lo et al., 2009). Thus, without reliable reproductive protocols for generating timed pregnant *Peromyscus*, most established forms of genetic engineering, including modern targeted integration methods based on CRISPR, cannot advance.

In this work, we detail our efforts to generate highly efficient pregnancy and pseudopregnancy events in *Peromyscus* in order to enable genome engineering in this and other non-model rodent species. We focused our attention on engineering *Peromyscus leucopus*, the primary reservoir of *Borrelia burgdorferi*, as engineering advances in this species may lead to new ecological tools for combating Lyme disease (Buchthal et al., 2019). We first attempted to derive stem cells from *P. leucopus* blastocysts and fibroblasts, which were subsequently engineered with CRISPR, but the cultures could not be reliably expanded. We next attempted i-GONAD for *Peromyscus* embryo engineering, but, when coupled with pregnancy block, our inefficient mating strategy resulted in low pup yield and even lower editing efficiency. We consequently refocused our efforts on developing a broadly enabling solution for improving mating efficiency using implanted temperature sensors to detect estrous-dependent changes in body temperature. In so doing, we identified that animal movement patterns are equally indicative of estrous cycle stage, thereby enabling the detection of ovulation (rather than simply mating events) via low-cost camera-based activity tracking. With reliable estrous monitoring, we demonstrate that the generation of pregnant and pseudopregnant *Peromyscus* is achievable with rates nearing *Mus*, enabling targeted transgenesis with CRISPR/Cas9. Finally, we demonstrate our method’s versatility across highly divergent species by successfully tracking the estrous cycles of both *Mus* and Syrian hamsters, an achievement that paves the way for efficient reproductive protocols and transgenesis in many more non-model rodent species.

## RESULTS

### Efforts to derive *Peromyscus* ESCs and iPSCs

To achieve transgenesis in *Peromyscus*, we began by attempting to derive *Peromyscus* embryonic stem cells (pESCs) from blastocysts. To generate fertilized *Peromyscus* embryos for ESC derivation, we attempted an optimized superovulation protocol provided by the Fisher lab (**Supp Fig. 1A**). However, hormone priming proved inefficient, resulting in a paucity of embryos. During each harvest, embryos were recovered at all stages of embryogenesis, including at the 1-to 4-cell stage (**Supp Fig. 1B**). The highest rates of development to the blastocyst stage *in vitro* were achieved by harvesting at the 8-cell stage. Five rounds of pESC derivation were attempted using embryos harvested at this stage (**Supp Fig. 1C**). The first round of pESC derivation was performed using mESCD-KSR medium (Markoulaki et al., 2008). A single outgrowth formed, which was dissociated and transferred to another well containing MEFs and mESCD-KSR. Several days after dissociation, small pESC-like colonies were observed. Like mESCs, pESC-like colonies were tightly formed, rounded and bright under phase contrast. However, there was a high rate of spontaneous differentiation among these pESC-like colonies, which failed to reliably expand and disappeared after several passages. To improve pESC derivation, we experimented with several published media formulations, including mESCD-KSR-2i, both with and without N2B27 (**Supp Fig. 1C**), which was previously found to facilitate ESC derivation in rats (Buehr et al., 2008). In our fifth attempt, we plated 9 blastocysts in mESCD-KSR-2i. All formed large outgrowths, and five gave rise to ESC-like colonies that successfully propagated. We assessed the pluripotency of these pESCs lines by staining for ESC pluripotency markers (**Supp Fig. 1D**) and by performing a teratoma formation assay, from which we obtained evidence of pESC contribution in the mesoderm and ectoderm (**Supp Fig. 1E**). However, we consistently encountered issues with pESC expansion and differentiation.

We simultaneously attempted to derive induced pluripotent stem cells (iPSCs). Briefly, we obtained mid-gestational fetal fibroblasts from *Peromyscus* fetuses (**Supp Fig. 1F**) and infected them with a reprogramming cocktail containing human cDNAs for KLF4, OCT4, SOX2, and c-MYC under the control of the tetracycline operator (Soldner et al., 2009). In the presence of doxycycline, we observed colonies with an apparent iPSC morphology. However, colonies differentiated upon doxycycline withdrawal and dox-independent clones were not obtained.

### Efforts to engineer *Peromyscus* ESCs

We next attempted to use our previously derived pESC lines to evaluate CRISPR-based engineering by encoding GFP in the *Peromyscus* Rosa26 locus. To identify the *Peromyscus* Rosa26 homology arms and corresponding CRISPR gRNA, we *de novo* sequenced the genome of our *Peromyscus leucopus* colony and compared it to another publicly available *P. leucopus* genome and other rodents (Long et al., 2019). The orthologous *Peromyscus* locus was identified using previously published Rosa26 *Mus* knock-ins (Chu et al., 2016)(**Supp Fig. 2A**). Two GFP+ pESC cell lines were generated following nucleofection of Cas9 + gRNA (**Supp Fig. 2B**), with varying success. Cells targeted with our first construct (pR26-CMV-EGFP) resulted in mixed populations of EGFP positive and negative cells due to challenges with dissociation and single cell expansion (**Supp Fig. 2C**). To enable selection and enhance fluorescence, we created a second targeting construct, pR26-CAGGS-EGFP-2A-Puro, with a selectable marker and a more efficient promoter which resulted in a homogeneous population of brighter EGFP positive cells (**Supp Fig. 2C**). These pESCs remained puromycin-resistant and GFP positive after multiple passages with selective media. However, when karyotyped, pESCs exhibited trisomies for chromosomes 16 & 17 (**Supp Fig. 2D**). Due to abnormal karyotyping and the aforementioned challenges with pESC propagation and expansion (see Supp. methods), efforts to further develop pESC-based strategies for genetic engineering were discontinued. While our iPSC-based efforts are ongoing, the difficulty of generating blastocysts led us to seek alternative methods.

### *In situ* germline genome engineering via i-GONAD

We next attempted to directly engineer *Peromyscus* embryos using i-GONAD, an electroporation method for generating germline-genome engineered animals which has been successfully implemented in multiple rodent species (**Fig. 1A**) (Ohtsuka et al., 2018). First, we validated the intraoviductal injection and electroporation technique by delivering EGFP mRNA into preimplantation embryos via *in situ* electroporation. Fluorescence was observed in recovered embryos, confirming successful mRNA delivery via i-GONAD (**Supp Fig. 2E**). We next attempted to engineer *Peromyscus* embryos with i-GONAD by delivering CRISPR ribonucleoproteins, including our previously validated *Peromyscus* Rosa26 CRISPR guide, into the oviducts of female *Peromyscus* via *in situ* electroporation. We performed 39 i-GONAD procedures on previously mated female *Peromyscus*, which gave rise to 4 litters, only two of which survived. Sequence analysis showed that one of the two litters contained a single gene-edited pup with a mutation in the *Peromyscus* Rosa26 locus (**Supp Fig. 2F**).

**Figure 1:**
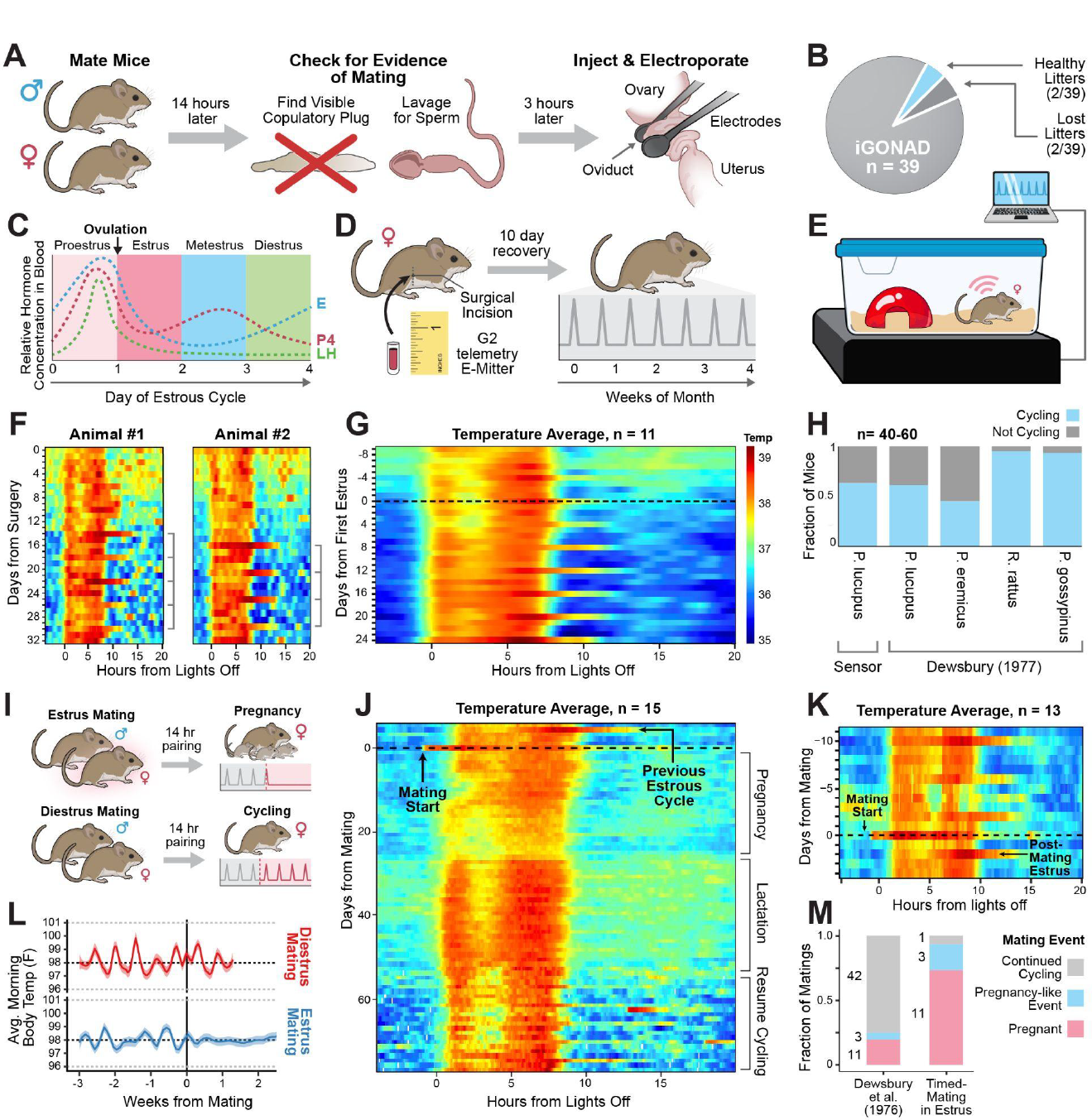
**A)** Diagram of iGONAD: Mice are mated and then checked for signs of pregnancy. CRISPR/Cas9 is injected into the oviduct of a pregnant mouse, and then the oviduct is electroporated. **B)** iGONAD statistics including all procedures performed in *Peromyscus*. **C)** The *Mus musculus* estrous cycle. Ovulation occurs every 4 to 5 days. **D)** Diagram of mouse telemetry system: G2 E-Mitter is implanted in the abdominal cavity in order to detect core body temperature and activity data over time. **E)** The G2 E-Mitter transmits data at 1 min intervals to a receiver, which is positioned under an animal’s cage and communicates with the Vital View Telemetry Software. **F)** Core body temperature data from two individual female *Peromyscus* with sequential days stacked from top to bottom. Prolonged peaks in core body temperature (estrus) appear in 4 day intervals. X axis: 0 = 8pm when lights turn off in the animal facility. Y axis: 0 = first day after surgery. **G)** Mean core body temperature for 11 female *Peromyscus* shows no variability in cycle length among cycling mice. **H)** Estrous cycling rate among all *Peromyscus* with implanted G2 E-Mitters compared to previously published cycling rates. **I)** Diagram of two different *Peromyscus* mating schemes. Top: Mating when mice are in estrus leads to the suspension of the estrous cycle. Bottom: When mice are mated in diestrus, the estrous cycle continues unabated. **J)** Mean core body temperature for 15 female *Peromyscus* mated in estrus. Sequential days are stacked from top to bottom from time of mating with stud males. Estrous cycle disruption is clearly visible after mating, indicating a high rate of pregnancy and pregnancy-like events. **K)** Mean core body temperature for 13 female *Peromyscus* mated in diestrus. All mice continued to cycle after mating. **L)** When compared, morning core body temperature for pregnant mice is distinct from non-pregnant mice, which show no disruption of morning temperature cycling when mated in diestrus. X axis: average core body temperature in the morning (after lights turn on). Y axis: 0 = mating in diestrus or estrus. **M)** Efficiency of timed pregnancy using core body temperature for estrous tracking compared with previously reported timed pregnancy rates in *Peromyscus* using vaginal lavage and cytology.

While we successfully employed i-GONAD to engineer *Peromyscus* deletions, the method proved to be highly inefficient; yielding only one gene-edited pup after nearly 40 surgeries (**Fig. 1B**). This poor efficiency may be partially attributed to a low rate of fertilization among the females subjected to i-GONAD, since vaginal lavage was used to detect sperm, which is an unreliable pregnancy marker in *Peromyscus leucopus (Dewsbury & Lanier, 1976)*. Moreover, pregnant *Peromyscus* that were repeatedly handled may have experienced pregnancy block, a phenomenon which has been reported to occur at a high rate in experimentally manipulated pregnant *Peromyscus* (Utt, 1984). *Collectively, these efforts underscored the need for a more efficient approach to generating timed pregnant Peromyscus*, which are required for nearly every transgenesis method.

### Estrous tracking with core body temperature

Timed mating is required for a variety of transgenesis protocols, including the generation of 1) embryos at a precise gestational age, 2) recipient mice suitable for embryo transfer, or 3) to implement iPSCs/ESC-based transgenesis. In *Mus*, achieving efficient pregnancy requires either properly timing the estrous state of female mice with mating to ensure receptivity to male studs, and/or checking females for vaginal plugs to determine if mating has occurred (Byers et al., 2012). The latter technique cannot be used with *Peromyscus* because their vaginal plugs are internal and undetectable (Hartung & Dewsbury, 1978). Researchers have previously employed vaginal cytology in *Peromyscus* to determine the stage of a mouse’s estrous cycle, which is reported to span 4 to 5 days, but with high variability in the length (ranging from 2 to 8+ days) (Dewsbury et al., 1977). When coupled with low timed pregnancy rates among *P. leucopus* subjected to vaginal lavage, existing estrous detection methods are demonstrably inadequate. Over the course of the estrous cycle, mice experience numerous physiological changes, including changes in body temperature, which result from hormonal changes during the time of ovulation (Marrone et al., 1976; Weinert et al., 2004). In particular, the release of progesterone, which is critical for the establishment and maintenance of pregnancy, causes a corresponding rise in body temperature during estrus (**Fig. 1C**). As a result, core body temperature telemetry systems have been successfully employed in *Mus musculus* to resolve the estrous cycle for individual laboratory mice and arrange highly efficient timed matings (Smarr et al., 2016). Core body temperature data collected by researchers using the aforementioned telemetry system supports a 4 to 5 day estrous cycle in *Mus*, as previously established *(Byers et* al., 2012). Estrus can be clearly observed in mice with implanted temperature sensors as prolonged heat periods that extend into the daily rest period. We sought to apply the same telemetry system to *Peromyscus* to resolve the estrous cycle and precisely ascertain when individual mice are cycling. Briefly, sensors (G2 E-Mitter, Starr Life Sciences, Oakmont, PA) were implanted in the abdominal cavity of 10 to 16 week old female *P. leucopus*. These sensors track core body temperature and locomotor activity when coupled with a receiver that powers the sensor and collects measurement data over time (**Fig. 1D**). Temperature readings were collected once per minute during the study (**Fig. 1E**).

Following temperature sensor implantation, cycling patterns were observed among *Peromyscus* from our colony beginning 3 to 14 days after surgery (**Supp Fig. 3**). Without exception, a clear 4 day estrous cycle was observed in each cycling mouse, manifesting as a peak in core body temperature after the lights were turned on (**Fig. 1F**). This trend was apparent for both individual mice and for the population average (**Fig. 1G**). Contrary to previous findings, no variability in the length of estrous cycle was observed; if a mouse began cycling, it cycled regularly every 4 days. In total, 57 temperature sensor implant surgeries were performed and 36 cycling mice were identified (**Supp Fig. 4**). The rate of cycling across all experiments (63%) was consistent with the reported rate of cycling among *P. leucopus* in a previous study (61%) (**Fig. 1H**) (Dewsbury et al., 1977).

### Pregnancy with estrous tracking

We next sought to establish whether mating *Peromyscus* in estrus, as determined by core body temperature, could boost the pregnancy rate from timed mating in this species. First, we monitored 15 female *Peromyscus* to determine the timing of their estrous cycles and then mated them with proven studs on the night of presumed estrus (**Fig. 1I**). After 4 days, suspended estrous cycles were observed in 14 out of 15 mice (**Supp Fig. 5**). The high rate of estrous cycle disruption is readily apparent on the population average (**Fig. 1J**). 11 of the 14 *Peromyscus* produced litters after 24 to 26 days, with an average litter size of 3.4 pups (**Supp Fig. 6**). Importantly, when female mice were mated during diestrus, no disruption of the estrous cycle was observed (**Fig. 1K**) and periodic heat fluctuations continue unabated unlike estrus matings (**Fig. 1L**). Cycling disruption, therefore, appears to be indicative of pregnancy, which is contingent on timed mating in estrus.

Interestingly, the telemetry system also revealed a subset of mice that experienced pregnancy-like events; 3 mice exhibited a break in estrous cycling that lasted for at least 20 days (**Supp Fig. 5**). Such pregnancy-like events have been previously documented in *Mus* by researchers employing mouse telemetry to improve timed pregnancy (Smarr et al., 2016). However, without further study, we cannot be sure if the observed suspension of the estrous cycle is due to implantation failure, pseudopregnancy or miscarriage. Overall, mouse telemetry for estrous detection more than quadrupled the efficiency of pregnancy or pregnancy-like events in *P. leucopus* from 20% to 93% (**Fig. 1M**).

### Estrous tracking with animal activity

As with core body temperature, researchers have observed elevated activity on days of estrus (Smarr et al., 2016). However, tracking mouse activity was previously deemed unreliable for estrous detection, since researchers could only distinguish estrus from averaged activity data, and not for individual mice (Smarr et al., 2016). In light of these findings, we had initially focused our efforts on resolving the *Peromyscus* estrous cycle from core body temperature readings alone, with a goal of ultimately detecting pregnancy or pregnancy-like events. However, once temperature-based estrous detection was established, we observed that temperature readouts were highly correlated with concurrent activity data collected by the sensors (**Supp Fig. 7A**). Notably, we were able to clearly identify a prolonged extension of activity coinciding with elevated temperature in individual mice on days of estrus (**Supp Fig. 7B**), a trend that was similarly visible at the population level (**Supp Fig. 7C**). Based on these findings, monitoring locomotor activity appears to be equally effective for establishing estrous.

While the mouse telemetry system enabled efficient timed mating in *Peromyscus*, it comes with significant drawbacks. First and foremost, each mouse must undergo an invasive sensor implantation surgery. Although surgery does not prevent cycling, it clearly disrupts the *Peromyscus* estrous cycle, as witnessed by the extended period between surgery and the first day of cycling, which spanned on average 13 days. Undoubtedly, the combined effects of post-surgical recovery and estrous disruption extend experimental timelines. Furthermore, this mouse telemetry system is expensive, with costs exceeding $1,575 per sensor set-up. This price point likely renders sensor-based estrous tracking infeasible for labs and transgenic cores to adopt at scale. As such, we next sought to develop a new estrous tracking system based on activity detection that would reduce both the cost and experimental delays precluding the application of highly efficient timed matings to entire colonies.

### Development of camera-based activity tracking for estrous detection

Unlike mouse telemetry, camera technology is cheap, widely available and noninvasive. Thus, we developed a camera system for tracking *Peromyscus* activity using the Raspberry Pi, a tiny, single-board computer that has been widely adopted due to its low cost, modularity, and open design. Briefly, *Peromyscus* cages were equipped with a Raspberry Pi Zero 2 W (WiFi-enabled) computer attached to a Raspberry Pi NoIR V2 camera module capable of medium-resolution photographs (**Fig. 2A**). Given the small size of the camera set-up, we eventually outfitted three mouse racks with 210 Raspberry Pi units to monitor the maximum number of mouse cages that can fit per rack (**Fig. 2B**). The camera module captured images at 4 second intervals, which were then processed on the Raspberry Pi units and the resultant activity data uploaded to our cloud database. Every two hours, this data was downloaded to the server, processed further, and cached for fast visualization. Every six hours, email notifications were sent out detailing any Pi units that hadn’t successfully uploaded their data (**Fig. 2C**). We created a web interface to view activity patterns and simplify equipment monitoring and troubleshooting (**Fig. 2D**).

**Figure 2:**
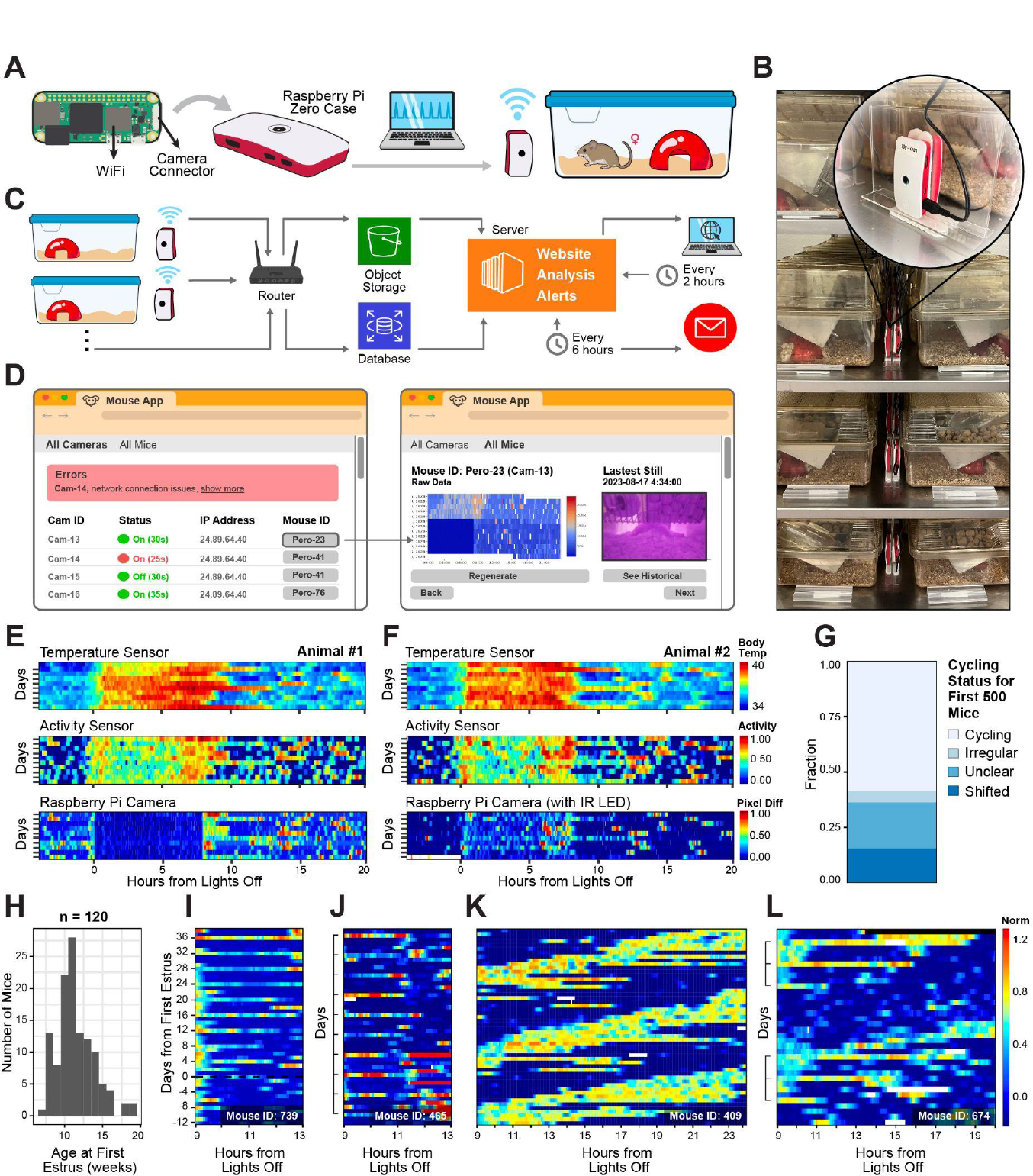
**A)** Diagram of camera system for tracking *Peromyscus* activity. Mouse movement was detected with a Raspberry Pi NoIR Camera V2 module attached to a Raspberry Pi Zero 2 W computer with WiFi. **B)** Raspberry Pi units were attached to acrylic stands that were positioned behind every cage on the cage rack. **C)** Diagram of data export and analysis. Images captured by Raspberry Pi cameras were processed on the Raspberry Pi units. The activity data was uploaded to the cloud database and then downloaded to the server and displayed on the website. **D)** Diagram of web interface. Active Raspberry Pi units were listed on the “All Cameras” page, along with their status and IP address, for equipment monitoring and troubleshooting. All mice were listed on the “All Mice” page with a click-through to individual mice, their activity graphs and latest photo to monitor for obstructions. **E)** Comparison of mouse estrous tracking systems for a single mouse using either core body temperature or activity from the mouse telemetry system, or activity from a Raspberry Pi unit. **F)** Comparison of all three estrous tracking systems on a different mouse, with a near-IR LED attached to the Raspberry Pi unit for night-time tracking. **G)** Cycling status of the first 500 mice placed on camera. **H)** Distribution of ages at which estrus activity was first detected. **I)** Example of early estrous onset. The first estrus behavior can be distinguished from non-estrous activity when no discernible 4-day trends are present. X axis: 0 = 8pm when lights turn off in the animal facility. Y axis: 0 = first estrus day. **J)** Rare example of *Peromyscus* with 5-day estrous cycle. **K)** Example of mouse with shifted circadian rhythm. **L)** Example of mouse with shifted circadian rhythm and visible estrous cycling behavior.

The G2 E-Mitter sensors detect movement through a voltage shift in an internal capacitor. Consequently, the act of scratching could register as activity as readily as lateral cage movements. Thus, we first sought to establish if estrous cycling activity was detectable via camera by placing a Raspberry Pi camera adjacent to a cage with a mouse with an implanted sensor. In doing so, we were able to simultaneously track both the core body temperature and locomotor activity data generated by the sensor, and any movement visible to the camera (**Fig. 2E**). Estrous cycling was easily identifiable by all three readouts, with estrus days as recognizable with the Raspberry Pi camera system as with telemetry. We further observed that the addition of a near-IR LED provided comparable resolution during the daytime while also enabling night-time tracking (**Fig. 2F**). In contrast to our mouse telemetry system, the introduction of cameras allowed us to observe cycling trends without the delays and discomfort caused by surgery. Furthermore, the cost of the camera setup is $75 per cage, which is less than 1/20 of the cost of mouse telemetry. Thus, our mouse camera system represents a cheap and effective alternative to invasive telemetry systems and can be readily implemented by any lab.

### Insights from camera-based activity tracking

As our colony grew, we eventually placed more than a thousand female mice under camera surveillance. The estrous cycling rate in our colony detected by camera mirrored the rate initially detected by mouse telemetry, confirming the effectiveness of our new approach (**Fig. 2G**). Female *Peromyscus* are believed to attain sexual maturity between 46 and 51 days of age (Clark, 1938). Yet, by monitoring a substantial cohort of prepubescent mice through sexual maturity, we identified an overall later onset of estrus behavior (mean = 11.5 weeks, sd = 2.5 weeks) and a broader spectrum of potential estrus initiation ages (**Fig. 2H**). The first estrus behavior stands out distinctly from non-estrus activity when no discernible trends are present (**Fig. 2I**) and almost always exhibits a consistent 4-day cycle. Out of over 1000 female *Peromyscus*, fewer than 5 mice have been observed with 5-day cycles (**Fig. 2J**).

Historically, *Peromyscus* are not associated with sleep disorders or any irregularities in their sleep patterns. However, camera-based activity tracking also revealed that approximately 20% of our mice displayed shifted circadian rhythms (**Fig. 2K**). It remains uncertain whether this phenomenon is unique to our colony, a result of inbreeding, or indicative of a broader trend in the population. This altered sleep pattern became evident soon after mice were placed under camera surveillance and remained consistent; it neither developed gradually nor dissipated over time. In some cases, estrous cycling could still be observed among mice with shifted circadian rhythms (**Fig. 2L**).

### Timed mating for embryo production

Having successfully implemented camera-based estrous tracking, we next sought to employ timed mating at scale for embryo generation. As previously mentioned, our superovulation protocol, which required a 72-hour gap between hormone administrations, yielded poor results (**Fig. 3A**). Following superovulation, embryos from a group of mice were harvested and pooled. Despite many efforts, the average yield per mouse remained below five embryos, and displayed marked inconsistency, with one experiment yielding fewer than one embryo per mouse (**Fig. 3B**). To identify the source of variability, we began tracking the number of cells harvested from individual mice (rather than bulk harvesting). We quickly discovered that a small group of mice was producing the vast majority of cells (**Fig. 3C**), leading to experimental inconsistency. Additionally, our fertilization rate was far lower than anticipated; even if a mouse produced eggs after superovulation and mating, they were unlikely to be fertilized (**Fig. 3D**). Following the identification of a clear 4-day estrous cycle, we implemented the standard *Mus* superovulation regimen of 48 hours between hormone doses, but unfortunately observed equally low rates of fertilization (**Fig. 3D**). Researchers have demonstrated that mouse oocytes and embryos produced without superovulation are of a superior quality to those produced after the exogenous administration of gonadotrophins (Ertzeid & Storeng, 2001; Lee et al., 2017). Given our high pregnancy efficiency with timed mating, coupled with the known embryo quality improvements, we next compared timed mating and superovulation. Briefly, mice were tracked via camera monitoring, and then mated on the night of presumed estrus. Embryos were then harvested and counted per mouse. This approach yielded far more uniform results; the variation in cells per animal diminished greatly in comparison to superovulation (**Fig. 3E**), while the number of fertilized embryos per mouse surged (natural mating mode embryos/mouse = 5, superovulation mode embryos/mouse = 0). Timed mating was highly efficient at embryo generation; when mice were paired during estrus, they consistently generated eggs and the majority were fertilized (**Fig. 3F**). Younger timed mated mice had greater embryo yields and fewer unfertilized eggs, whereas older mice displayed reduced productivity (**Fig. 3G**). Given the observed inconsistencies, lower yield and fertilization rate with superovulation, coupled with the superior quality of embryos obtained through natural mating, the latter appears to be a more reliable and effective method for embryo production in *P. leucopus*.

**Figure 3:**
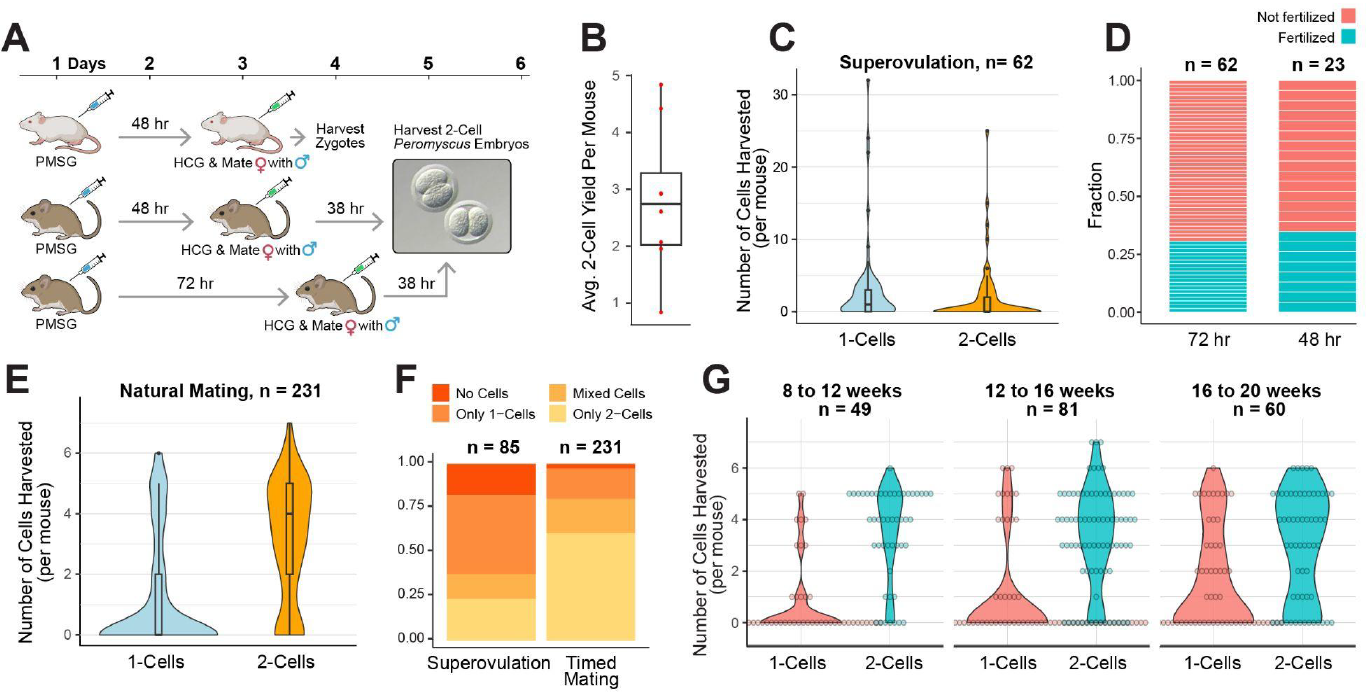
**A)** Diagram of *Mus* and *Peromyscus* hormone priming protocols. **B)** Pooled results from superovulation experiments, in which mice were hormone primed with 72 hours between PMSG and HCG. The average embryo yield per mouse was highly variable between experiments. **C)** Cells harvested from individual mice following the 72 hour superovulation protocol, demonstrate low numbers of both 1 and 2 cell embryos, with outliers. **D)** Comparison of fertilization rate following either the 72 hr or 48 hr superovulation protocols; both produced equally low rates of fertilized cells. **E)** Cells harvested from individual mice after timed mating without superovulation. Timed mating resulted in more fertilized embryos per mouse and less variability in cell yield. **F)** Fraction of cell types harvested from superovulated or timed mated mice. **G)** Cells harvested from timed mated mice segmented by age. Younger mice produced higher 2-cell embryo yields than older mice, which were more likely to produce unfertilized eggs.

### Pseudopregnancy generation with estrous tracking

Most transgenesis efforts require the efficient generation of pseudopregnant recipients through the identification of copulatory plugs, which are not readily observable in *P. leucopus*. Thus, we sought to use estrous tracking to enable the efficient generation of pseudopregnant mice. Briefly, male *Peromyscus* were vasectomized and sterility confirmed through mating experiments. Next, 6 female *Peromyscus* were tracked with implanted temperature sensors. Once estrous was established, females were mated on the night of presumed estrus with vasectomized males (**Fig. 4A**). After 4 days, suspended estrous cycles were observed in 5 of the 6 mice. The high rate of estrous cycle disruption is apparent among the population average (**Fig. 4B**). All mice resumed cycling roughly 2 weeks later (range = 10 to 14 days) (**Supp Fig. 4**). When core body temperatures are compared, pseudopregnancy closely resembles pregnancy in *Peromyscus*, except in length (**Supp Fig. 8**). A total of 54 camera-tracked female *Peromyscus* were mated with vasectomized males and then subjected to various interventions for the experiments discussed in the subsequent sections. Of the 53 mice with data post mating, 85% (45/53) had suspended estrous cycling (**Fig. 4C**). Interestingly, for 6 of the 8 mice that continued to cycle, their cycle shifted by one day post mating (**Supp Fig. 9**).

**Figure 4:**
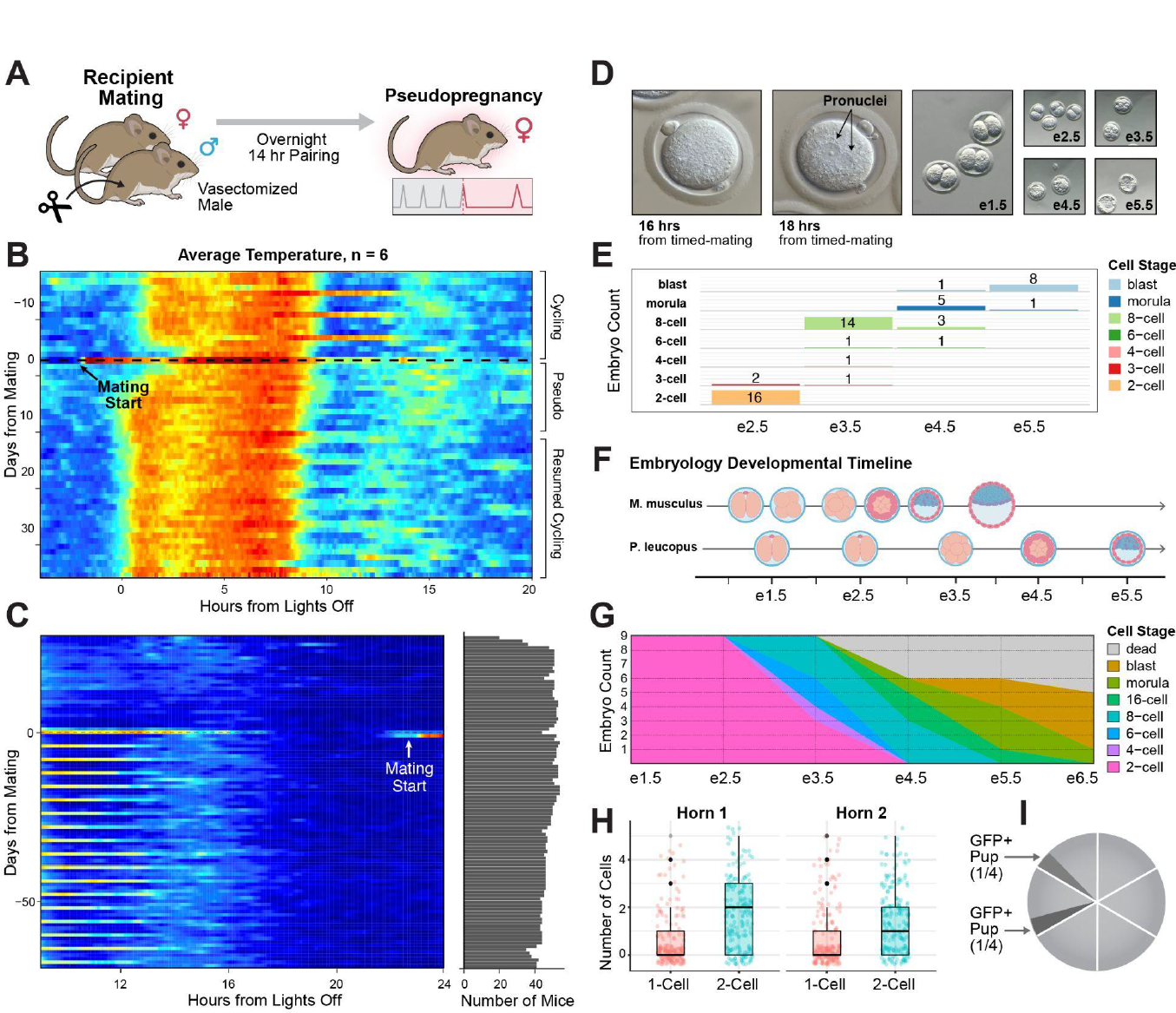
**A)** Diagram of mating scheme for pseudopregnancy in *Peromyscus*. Mating in estrus with a vasectomized male is expected to produce pseudopregnant females with suspended cycles. **B)** Mean core body temperature for 6 mice with implanted sensors that had been mated in estrus with vasectomized males. Estrous cycle suspension is visible after mating, indicating a high rate of pseudopregnancy. X axis: 0 = 8pm when lights turn off in the animal facility. Y axis: 0 = mating day. **C)** Left: Mean activity plot for 53 camera-tracked mice mated with vas males in estrus. 85% of mice exhibited suspended estrous cycling. Right: Number of mice per time point. **D)** Photographs of early gestational time points, including zygote development, from timed mated mice. The pronuclei only became visible 18 hours after timed matings were set up. **E)** Distribution of cells harvested on gestational days e2.5 through e5.5 from timed mated mice. **F)** Diagram of Mus and *Peromyscus leucopus* early gestational timelines. Peromyscus embryos stay in the 2-cell stage longer than *Mus* embryos and require considerably more time to reach the blastocyst stage. **G)** In vitro developmental timeline of *Peromyscus* embryos harvested at the 2-cell stage and cultured in KSOM until blastocysts formed. **H)** Cells recovered from individual uterine horns from timed mated mice on gestation day e1.5. **I)** Pronuclear injection statistics for the 6 recipients that each received 4 embryos injected with Cas9, Rosa26 guide and the EGFP targeting construct.

### *P. Leucopus* developmental timelines *in vivo & in vitro*

While the developmental timeline for *P. maniculatus* is well-documented, there is scarce information on the early developmental stages crucial for implementing embryology protocols in *P. leucopus* (Davis & Keisler, 2016). *Further, these established timelines are impacted by inherent variability stemming from the lack of reliable timed matings. Thus, we sought to establish a comprehensive developmental timeline for Peromyscus leucopus* in order to determine the optimal timings for embryo harvest and transfer. We performed 40 timed matings using camera-tracked female mice. We recovered embryos for daily early gestational time points (**Fig. 4D**) and throughput development. Pronuclei became visible 13 hours post-mating (or after 18 hours in total), assuming fertilization occurs at the midpoint of the dark cycle. In contrast, the *Mus* pronucleus appears 5 to 11 hours after fertilization. For the early gestational stages, we mapped the cell types retrieved at each time point (**Fig. 4E**). Compared to *Mus musculus, Peromyscus* embryos exhibit extended two-cell stages, persisting until e2.5, and demonstrate delayed blastocyst development, with blastocysts forming on day e5.5 (**Fig. 4F**). Blastocyst implantation appears to occur on or before e6.5 as no cells were found at this time point. The extended two-cell phase in *Peromyscus* could potentially increase the efficiency of genomic editing, given a potentially higher rate of homologous recombination during a longer G2 phase (Gu et al., 2018). Since transgenesis protocols often necessitate extended overnight embryo culture, we next sought to characterize the *in vitro* development of *Peromyscus* embryos. Harvested 2-cell embryos from timed mated mice were cultured in KSOM and housed in a tri-gas incubator until blastocysts formed. Embryos were imaged daily in order to construct an *in vitro* developmental timeline (**Fig. 4G**). We observed that 44% of the embryos developed to the blastocyst stage, and that development was delayed *in vitro*, with blastocysts appearing on day e6.5. Given these findings, we chose to forgo the standard overnight culture protocol to avoid developmental delays, pending further refinement of culture conditions.

### *P. Leucopus* embryo transfer & pronuclear injection

In preparation for embryo transfer, we sought to identify the ideal number of embryos to transfer by studying the inherent capacity of *Peromyscus* oviducts. We mated camera-tracked female *Peromyscus* on the night of presumed estrus, then harvested embryos on gestation day e1.5, recording quantities from each individual uterine horn (**Fig. 4H**). Noting a maximum of five embryos per uterine horn, we concluded that transferring four embryos would have the greatest potential for successful implantation.We transferred unedited embryos into pseudopregnant *Peromyscus* to establish a baseline implantation rate, with the eventual goal of transferring engineered embryos. Briefly, we organized two groups of female mice, embryo donors and embryo recipients, and established their estrous cycles with camera-based tracking. To synchronize reproductive stages, donors were mated with proven studs a day before recipient matings with vasectomized males. This scheduling allowed for the harvest and implantation of two-cell embryos on the same day, eliminating the need for overnight incubation. Embryos were harvested on gestation day e1.5 in M2 and underwent brief culturing in KSOM within our tri-gas incubator. Finally, four two-cell embryos were transferred into one oviduct for each recipient. In total, 12 embryo transfers of unedited *Peromycsus* cells were performed. 6 recipients produced a total of 15 pups or midgestation fetuses, as our embryo transfer efficiency improved over time (from 2/6 to 4/6 recipients). These findings demonstrate the viability of embryo transfer in *Peromyscus*, thereby enabling rederivation.

With embryo transfer established, we next sought to demonstrate genome engineering by performing pronuclear injection with CRISPR/Cas9. We generated a modified version of our second targeting construct in which the selection marker was removed (pR26-CAGGS-EGFP) (**Supp Fig. 2B**). Camera-tracked donors were mated with proven studs one day prior to the mating of camera-tracked recipients with vasectomized males in order to synchronize reproductive stages. Two-cell embryos (e1.5) were harvested, and each blastomere was injected with the targeting construct, along with Cas9 and guide RNA. A total of 24 CRISPR-injected two-cell embryos were transferred to six recipients (4 per mouse). Two pups were delivered, both of which were GFP-positive, resulting in an 8% conversion rate with 100% of delivered pups containing the insert (**Fig. 4I**). By comparison, previous attempts to engineer the *Mus* Rosa26 locus by 2-cell injection with a comparable insert (4.5kb), generated a total of two founders from twenty-two pups. following transfer of seventy-five microinjected embryos, for a 3% conversion rate with only 9% of pups containing the insert <u>(Gu et al. 2018)</u>. Thus, these advances demonstrate the practical application of CRISPR technology in generating transgenic *Peromyscus*, paving the way for genetic investigations in this species.

### Camera-based activity tracking of other rodent species

For over a century, researchers have attempted to link physical activity and estrous state in different species with mixed results (Slonaker, 1924; Smarr et al., 2016; Wollnik & Döhler, 1986; Wollnik & Turek, 1988). We next sought to determine whether camera-tracking could be used to identify periodic, cycling activity in other rodent species. As proof of concept, the camera-based tracking system was used to monitor female C57BL/6 *Mus musculus*, where estrous activity is reportedly undetectable at the individual level (Smarr et al., 2016), and Syrian hamsters *Mesocricetus auratus*, a species previously studied for activity patterns during their estrous cycles (**Fig. 5A**)(Prendergast et al., 2012). **For both *Mus* and hamsters**, behaviors associated with estrous cycling are reported to manifest at night (Prendergast et al., 2012; Smarr et al., 2016), in contrast to P. leucopus, which extends into the morning. Thus, having previously validated nighttime camera-based tracking with additional IR illumination, IR light strips (λ=940nm) were used to illuminate the cages. IR cameras were used to track a total of 14 female *Mus musculus* and 5 female hamsters. Shortly after the animals were placed on camera, a 5-day cycling pattern between the hours of 6 and 9 UTC became apparent for one of the *Mus* (**Fig. 5B**). However, these periodic activity trends were much harder to discern by eye for both *Mus* and hamsters. The camera-based activity tracking system detects activity via pixel-wise subtraction on two successive images (see supplemental methods). When images are taken under near infra-red illumination, several issues impede the detection of nocturnal animal activity. Under IR lighting, some animals become indistinguishable from the background (**Supp Fig. 10**). Further, confounding artifacts such as reflections on the cage walls and wire, shifting bedding, flickering IR light and intrinsic sensor noise, may be misinterpreted as movement. To improve activity detection, we applied background subtraction to images using the “TwoPoint” model (see supplemental methods), which classifies an image as either foreground or background. Additionally, each image was sectioned into quadrants, and the quadrant containing the food hamper was used exclusively for analysis. Following image processing, the one *Mus* with visible cyclical behavior showed periodic cumulative activity 5 to 8 hours after lights off; autocorrelation confirmed the significance of this observed 5-day trend (**Fig. 5C**).

**Figure 5:**
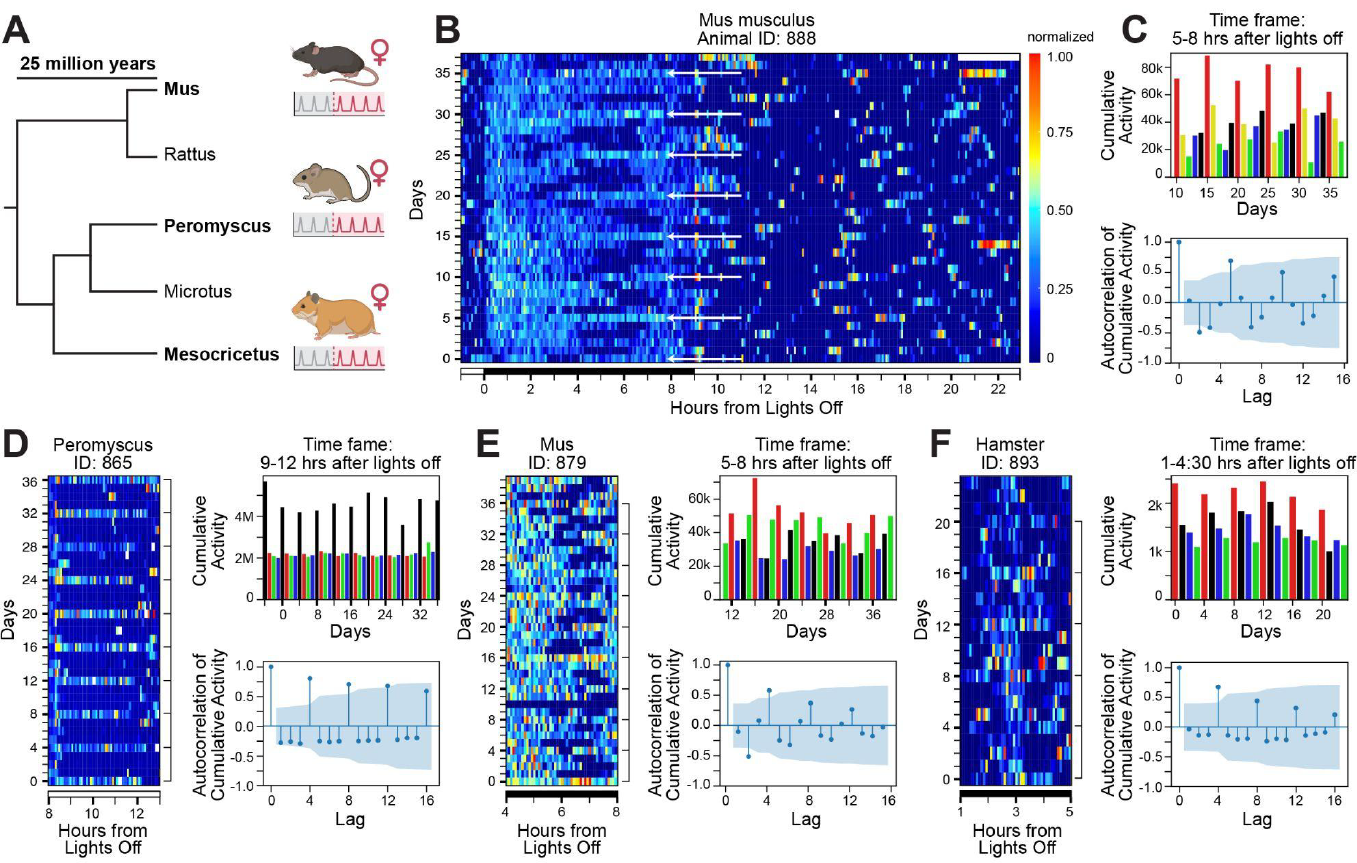
**A)** Phylogenetic tree illustrating the genetic distance between the genus *Mus* (from the Muridae family) and the genus *Peromyscus* and *Mesocricetus* or hamsters (from the Cricetidae family). Adapted from Bedford and Hoekstra (2015). **B)** *Mus* activity heatmap generated from image difference analysis showcasing a 5-day cycling pattern (indicated by white arrows) 5 to 8 hours after lights off. **C)** *Mus* daily cumulative activity plot with a 5-day color scheme (top) and its autocorrelation (bottom), post-sectioning and background removal; autocorrelation shown with a 95% confidence interval. **D)** *Peromyscus* activity heatmap (left), daily cumulative activity plot with a 4-day color scheme (top right) and autocorrelation with a 95% confidence interval (bottom right). All derived from image difference analysis without sectioning and background removal. **E)** *Mus* activity heatmap (left), daily cumulative activity plot with a 4-day color scheme (top right), and its autocorrelation (bottom right) post sectioning and background subtraction. **F)** Hamster activity heatmap detected by thermal camera (left), corresponding daily cumulative activity plot with a 4-day color scheme (top right) and its autocorrelation post sectioning and background subtraction with a 95% confidence interval (bottom right).

A significant positive spike at lag 5 and a negative value at lag 2 suggests an inverse relationship within this 5-day cycle, indicating alternating high and low activity days. By comparison, *Peromyscus* with clear periodic behavior 9 and 12 hours after lights off have much lower variability in daily cumulative activity. As such, autocorrelation was also statistically significant at lags 8 and 12 (**Fig. 5D**). Using only autocorrelation, periodic cycling behaviors were observed among a larger group of *Mus* and hamsters (**Supp Fig. 11**). While image processing did not greatly improve visual trend detection in *Mus*, daily cumulative activity revealed periodic 4 or 5-day patterns 5 to 8 hours after lights off (**Fig. 5E**), with varying significance (**Supp Fig. 11**). Interestingly, one Mus displayed a shift in its cycling day over time, a behavioral change rarely seen in Peromyscus without mating (**Supp Fig. 12**).

Periodic trends in hamsters were similarly not discernible by eye from activity graphs, but daily cumulative activity exhibited 4-day trends, with varying significance levels confirmed through autocorrelation (**Supp Fig. 11**). To improve activity tracking by distinguishing the animal more precisely from its environment, near-IR cameras were substituted with thermal cameras. Subsequently, hamster activity trends became visually apparent by activity graph 1 to 4.5 hours after lights off. While moving cage elements occasionally interfered with the signal, the periodicity of the 4-day cycling trend gained even greater statistical confidence (**Fig. 5F**). Collectively, these results suggest that simple, low-cost camera-tracking using near-IR or thermal cameras is a viable method for monitoring periodic behavior across a broad range of species, regardless of time of day or activity level.

## DISCUSSION

*Peromyscus* have been proposed as a model organism for human disease, as they appear to more closely resemble humans than laboratory mice by hematological measurements, methylation profiles, disease susceptibility, genetic diversity, and lifespan (which exceeds *Mus* in captivity by about threefold) (Sun 2014, Horvath 2022, Havighorst 2017). However, it is widely acknowledged that the use of *Peromyscus* as a disease model has been impeded by the dearth of genomic tools and technologies available for this species (Havighorst 2017). The challenges associated with engineering this species were and are formidable, as evidenced by the many issues we encountered when propagating and expanding ESCs, generating self-sustaining iPSCs, culturing *Peromyscus* embryos, hormone priming mice, and even simply maintaining a large enough colony to support our experiments without financial strain. Much work remains to be done in all of these areas.

Nevertheless, the advancements described herein mark the arrival of *Peromyscus* as a tractable model organism. We present highly optimized reproductive protocols for the generation of timed pregnant and pseudopregnant *Peromyscus* via low cost camera-based activity tracking, which rival the efficiency of timed matings in laboratory mice. Such protocols completely eliminate the need for plug checking, vaginal cytology and sperm detection, which are not only invasive, but known to cause pregnancy block at high rates in this species. Leveraging these protocols, we show that efficient embryo generation and transfer are achievable in *P. leucopus*, thereby unlocking the use of the most powerful transgenesis techniques, namely zygote and stem cell-based technologies, in this species. Finally, we show efficient *Peromyscus* genome engineering with CRISPR/Cas9, paving the way for the development of *Peromyscus* as homologous, isomorphic and predictive disease models, as well as their use for ecological engineering projects, such as Mice Against Ticks, a community-guided effort to engineer Lyme-resistant *Peromyscus* (Buchthal et al., 2019).

For more than a century, researchers have explored the relationship between physical activity and estrous states in various species, with varying degrees of success. A pioneering study published in the early 20th century was among the first to establish a link between rodent activity and estrous stages (Slonaker, 1924). However, subsequent studies challenged this correlation in mice and rats, contending that activity levels were not indicative of estrous cycling even in animals with previously established estrous cycles (Smarr et al., 2016; Wollnik & Döhler, 1986). Consequently, success in detecting estrous-linked activity appears to hinge on the monitoring technology applied.

To our knowledge, no researcher has utilized camera-based tracking for estrous detection, though it has become an essential tool for the study of animal behavior, providing unparalleled insight into behavioral development (Dunn et al., 2021), social learning (Hong et al., 2015), neuropsychiatric disorders (Woodard et al., 2017) and memory by tracing the locomotor activity of laboratory animals (Seibenhener & Wooten, 2015). Our efforts to apply camera-based tracking to a larger group of rodents has expanded its utility, by broadening the range of detectable movements and enabling monitoring regardless of the time of day. The identification of species-specific, periodic activity in three evolutionarily distinct rodents suggests that cycling behaviors are widespread.

With over 56 recognized species, the *Peromyscus* genus is celebrated for its extensive diversity, and the tools described herein are liable to unlock genome engineering for all *Peromyscus* species. Further, these advances hold promise beyond *Peromyscus*, and may extend the existing set of model organisms to include other intractable rodents, with major opportunities in less-studied systems for modeling human disease. Possible examples with translational potential include, but are not limited to, refractory species of rats, prairie voles, guinea pigs, and naked mole rats. By comparing well-established *Mus* strains to new rodent models, we may be able to use genetic engineering to dissect the underlying mechanisms that influence longevity (Holzenberger et al., 2003; Liang et al., 2003), neurogenesis (Kim et al., 2011), social behavior patterns (McGraw & Young, 2010), degeneration (Eimer & Vassar, 2013), and susceptibility to disease (Tian et al., 2013). Collectively, these tools for high efficiency timed pregnancy and pseudopregnancy represent a transformative opportunity for transgenesis in *Peromyscus* and broaden the use of genetic engineering technologies in recalcitrant species.

Finally, due to the abundance of wild *P. leucopus* in North America, and their role in the transmission of infectious disease, we have proposed immunizing local *P. leucopus* populations to be heritably resistant to *B. burgdorferi*, the etiological agent of Lyme disease (Iii et al., 2021). In doing so, we hope to break the disease transmission cycle between wild mice and ticks, and reduce disease prevalence in hard-hit areas like Nantucket and Martha’s Vineyard (Buchthal et al., 2019). By combining high-efficiency estrous detection with CRISPR/Cas9 genome engineering, we are poised to generate wild *P. leucopus* which heritability express anti-Lyme antibodies derived from locally sourced mice (Schaible et al., 1990). Engineered *Peromyscus* generated via camera-based estrous-tracking may serve as a proof of concept for engineering rodents to prevent zoonoses ranging from Lassa fever to hantavirus pulmonary syndrome.

## ACKNOWLEDGEMENTS

We offer our sincere thanks to Dr. Siniša Hrvatin for providing equipment for our preliminary estrous tracking experiments and for his guidance on sensor implantation and software configuration. We express deep gratitude to MIT’s Division of Comparative Medicine and its director, Dr. Kelly Metcalf Pate, for supporting our studies and for working to ensure that we preserve the privilege of conducting animal research. Additionally, we sincerely thank Dr. Susan Erdman, Dr. Bernard Varian, Julien Freeman, Andrea Vargas, Julie Anderson and Harlin Lemus, for their dedication to maintaining our *Peromyscus* colony and advancing our studies, and for their many thoughtful suggestions. We express our deepest appreciation to the communities of Nantucket and Martha’s Vineyard for their continued support and feedback, including Dr. Roberto Santamaria, Dr. Malcolm MacNab, Dr. Emily Goldstein Murphy, Dr. Jim Butterick, Dr. Carrie Fyler and Dr. Fyler’s many wonderful students. We offer our deep gratitude to Todd Rainwater and Amy Rommel for their enthusiasm, encouragement and thoughtful input. We are immensely grateful to the board of the Mice Against Ticks nonprofit, Ana Martinez and Beth Colt, for their commitment, counsel and invaluable guidance. We express our utmost appreciation to Adair Oesterle for crafting the finest needles on the planet, and for her invaluable guidance and friendship. We thank Dr. Heidi Fisher for generously sharing her lab’s superovulation protocol and for her kind support when we established our Peromyscus colony. We express gratitude to Tony Chavarria for his guidance in mouse surgery, many helpful suggestions and friendship. We thank Gabriel Meier for his assistance in adapting iGONAD to Peromyscus. We thank Domenick Giordano for his support during the set-up of our mouse telemetry system. We thank Dr. Alan Barbour for his assistance in the analysis of our *Peromyscus leucopus* genome and are grateful to Yelana (Helen) Skaletsky for her support in identifying the *Peromyscus* Rosa26 locus.

This manuscript is dedicated to the late Dr. Howard Dickler, a champion for the Mice Against Ticks project, whose invaluable guidance, boundless support, and exceptional warmth were ever-present and are deeply missed.

## AUTHOR CONTRIBUTIONS

### General Contributions

**JB:** Conceptualization, Formal Analysis, Resources, Writing-Original Draft, Visualization, Project Administration, Funding Acquisition; **EJC:** Conceptualization, Formal Analysis, Data Curation, Writing-Original Draft, Visualization, Supervision; **SRT**: Resources; **RJ**: Supervision; **SM:** Supervision; **KME:** Conceptualization, Methodology, Supervision, Funding Acquisition.

### Stem Cell Derivation/iGONAD

**JB**: Conceptualization, Formal Analysis, Funding Acquisition; **ZH**: Methodology, Investigation (ESC, iPSC, iGONAD); **SRT:** Resources; **RJ**: Conceptualization, Supervision; **SM:** Conceptualization, Methodology, Investigation (ESC, iPSC, iGONAD); **KME**: Conceptualization, Methodology, Resources, Funding Acquisition.

### Estrous Tracking & Transgenesis

**JB:** Conceptualization, Methodology, Validation, Formal Analysis, Investigation, Funding Acquisition; **EJC:** Conceptualization, Methodology, Software, Formal Analysis, Data Curation; **ZH:** Methodology, Validation, Investigation (Embryo Transfer, Pronuclear Injection); **CD:** Methodology, Software, Investigation, Data Curation (Camera Tracking); **BT:** Software, Formal Analysis, Investigation, Data Curation (Camera Tracking); **RPW**: Software, Investigation, Data Curation (Camera Tracking); **CC:** Investigation (Camera Tracking); **SG:** Methodology, Validation (Camera Tracking); **SRT:** Resources; **KLM:** Methodology, Resources (Camera Tracking); **KME**: Conceptualization, Methodology. All authors assisted in reviewing and editing the final draft of the manuscript.

## ETHICS STATEMENT

The Massachusetts Institute of Technology’s Committee on Animal Care approved all *Peromyscus* procedures and all *Peromyscus* were maintained at MIT (Cambridge, MA, USA) in strict accordance with all institutional protocols, DOD guidelines and the Guide for the Care and Use of Laboratory Animals. The Harvard University Institutional Animal Care & Use Committee approved all *Mus musculus* studies and all *Mus* were maintained at Harvard (Cambridge, MA, USA). The Tufts University Institutional Animal Care & Use Committee approved all hamster studies and all hamsters were maintained at Tufts (North Grafton, MA, USA).

## METHODS

### Superovulation of *Peromyscus*

Briefly, 4 to 6-week-old female *P. leucopus* were injected intraperitoneally with 20 IU of PMSG 1 hour prior to lights out followed by 20 IU of hCG 72 or 48 hours later. Mice were mated overnight and then embryos were harvested from donor females 36 hours after timed matings were set up.

### Mouse Embryonic Fibroblast Derivation (Feeders)

For MEF isolation, embryos were isolated at E13.5 from C57BL/6 mice and the fibroblast-containing tissue was collected under a dissection scope. The collected tissues were physically dissociated and incubated in Trypsin at 37 °C for 20 min after which cells were resuspended in MEF medium, γ-irradiated, and frozen for later use.

### pESC Derivation

For pESC derivation, 8-cell embryos were extracted at E3.0 from *Peromyscus* and plated in 96-well plates that had been coated with a mouse (*Mus*) embryonic fibroblast feeder layer. pESC derivation media conditions are detailed in Supp **Fig. 1C**. MEFs were pre-plated at least 12 h in advance in MEF medium. In preparation for the blastocyst plating, MEF medium was removed from the 96-well plates with feeders and replaced with 500 μl of pESC-derivation medium. Immediately prior to plating, the zona surrounding the blastocyst was removed by brief exposure to acid Tyrode’s solution. After zona removal, single blastocysts were placed in wells of 4-well plates, which were placed in a CO2 incubator and allowed to attach to the feeder layers. When an inner cell mass outgrowth was observed (5-6 days after plating), cells were first passaged using trypsin–EDTA. After a brief incubation at 37°C, cells were dissociated by pipetting up and down 3-4 times. After another short incubation, the contents of the wells were then transferred into a 12-well plate with feeders in the presence of pESC-derivation medium and further cultured.

### Attempt 1 details

In brief, using mESCD-KS the Zona Pellucida was removed from two blastocysts at E5.5 and embryos were plated individually on *Mus* embryonic fibroblast (MEF)-coated plates with mESCD-KSR.

### Attempt 2 details

including mESCD-KSR-2i, both with and without N2B27 (Supp **Fig. 1C**), next added 2i, a selective small molecule inhibitor of differentiation-inducing signals from GSK3β and ERK/MEK, to our mESCD-KSR medium. In our second attempt, only one blastocyst plated in the mESCD-KSR-2i condition formed an expanded outgrowth, which carried over, but never gave rise to a stable line. Next, we supplemented mESCD-KSR-2i with N2B27 which was previously found to facilitate ESC derivation in rats (Buehr et al., 2008) Our most promising results came from our final attempt: out of the 9 blastocysts plated in mESCD-KSR-2i, all we plated 9 blastocysts in mESCD-KSR-2i,, the same condition as our second attempt. All nine outgrowths formed large outgrowths and five gave rise to ESC-like colonies that became stable lines.

### pESC Dissociation and Propagation Challenges

Due to challenges associated with pESC propagation, our *Peromyscus* ESC-like colonies were not easy to maintain; grew slowly, differentiated frequently and were not ready to dissociate for 7-10 days after plating. Some colonies appeared to be resistant to dissociation, forming large clumps even after long incubations in Trypsin or Accutase. We employed a variety of dissociation reagents without detectable improvement, including higher concentration of Trypsin, higher concentration of EDTA in Trypsin, TryplE (GIBCO), TryplE Express, Collagenase, Pronase, Versin, Dispace, ReLeSR, as well as manual passaging. Moreover, MEFs appeared to be unhealthy, during the prolonged periods in culture between passages. Swapping MEFs inactivated by irradiation with Mitomycin C inactivated MEFs did not result in noticeable improvement. Using a lower density of MEFs alleviated some of the problems associated with higher density (including lifting off plates while trapping pESCs) but did not sufficiently support pESC propagation. We also tried culturing pESCs in the absence of MEFs by coating our plates with Matrigel or Gelatin, but most cells either didn’t attach or formed spheres/embryoid bodies that could not be dissociated. Due to aforementioned challenges, including spontaneous differentiation and issues with dissociation and inefficient propagation, further optimization of culture conditions is needed in order to facilitate pESC based transgenesis methods.

### pESC Staining

Cells were fixed in 4% paraformaldehyde for 20 min at 25 °C, washed three times with PBS, and blocked for 15 min with 5% FBS in PBS containing 0.1% Triton-X. After incubation with primary antibodies against Oct4 (Santa Cruz, h-134); Sox2 (R&D Biosystems); Nanog (anti-ms and anti-h, R&D Biosystems); for 1 h in 1% FBS in PBS containing 0.1% Triton-X, cells were washed three times with PBS and incubated with fluorophore-labeled appropriate secondary antibodies purchased from Jackson Immunoresearch. Cells were analyzed on an Olympus fluorescence microscope and images were acquired with a Zeiss Axiocam camera.

### Teratoma Formation

Teratoma formation was performed by depositing 1×10^6 cells under the flanks of recipient SCID mice. Tumors were isolated 3–6 weeks later for histological analysis. Mice were sacrificed before tumor size exceeded 1.5 cm in diameter.

### Generations of *Peromyscus* Tail Tip Fibroblasts

To generate primary fibroblasts for iPSC derivation, mice were sacrificed and a 1-2 mm piece tail was removed from 5-6 week old rodents. The tail tissue was then rinsed with Wescodyne (iodine-based antimicrobial solution) followed by 70% ethanol, and rinsed 2X in PBS with 1X Penstrep (Gibco). Working in a 6cm TC dish under a laminar flow hood, each tail biopsy was then finely minced with a sterile razor blade/scalpel into a 0.25ml mixture of collagenase and dispase neutral protease (4mg/ml each in DMEM). After mincing finely, 0.25 ml collagenase mixture was added to the plate, and incubated for 30 minutes at 37°C.

Following dissociation, 6mls of MEF media (DMEM +10%FBS + pen/strep) was added to the 6 cm plate and incubated overnight at 37°C, 5% CO2. The following day, to further aid dissociation of any large pieces of tissue, remaining tissue was titrated with a sterile Pasteur pipette. The tissue was then incubated for an additional 3 days at 37°C, 5% CO2 to allow cells to adhere and spread away from the tissue chunks. After 3-4 days, the media was refreshed, as cells began to undergo rapid proliferation. When nearly confluent, cells were passaged or frozen.

### iPSC Derivation

To create iPSCs, fully-differentiated adult somatic cells were returned to a pluripotent state using a cocktail of factors (hOct3/4, hSox2, hc-Myc, and hKlf4) that are known to maintain pluripotency during embryonic development. To generate *Peromyscus* iPSCs, the four factors, as well as the tetracycline transactivator, were delivered using tetracycline (Dox)-inducible lentiviral vectors. The lentiviral backbone for dox-inducible transgene expression was constructed by replacing the human ubiquitin C promoter of the FUW plasmid (Lois et al., 2002) with a tetracycline operator and minimal CMV promoter. For infections, virus was prepared by transfecting 293T cells with a mixture of viral plasmid and packaging constructs expressing the viral packaging functions and the VSV-G protein. Medium was replaced 24 hr after transfection, and viral supernatants were collected at 48 and 72 hrs. After filtration, supernatants were pooled, and tail tip fibroblasts were incubated with viral supernatants and fresh media at a ratio of 1:1 for 24 hr. They were subsequently cultured in ESC medium. Fully reprogrammed colonies were passaged for 12-20 days, at which point Doxycycline was withdrawn. Unfortunately, spontaneous differentiation was observed following removal of Dox, indicating lack of successful reprogramming.

### De Novo Sequencing of *Peromyscus* Genome

Blood samples were collected in a collection tube coated with an anticoagulant (EDTA) and inverted 8-10 times immediately following collection to ensure proper mixing of the anticoagulant. Blood was shipped on dry ice immediately following extraction to Dovetail Genomics (Santa Cruz) for *de novo* genome assembly.

### Guide Targeting Construct Design & Guide Design

Rosa26 sgRNA for iGONAD and pronuclear injection was purchased from Integrated DNA Technologies (Fig 2A). Rosa26 homology arms for ssDNA (for iGONAD) and targeting constructs (for pronuclear injection) were identified by aligning the *Mus* Rosa26 locus to the *Peromyscus* genome assembled via Dovetail Genomics. Homology arms were synthesized as gBlocks via IDT and either cloned into a standard high-copy number plasmid backbone, or used to make ssDNA, which was produced using the Guide-it Long ssDNA Production System v2 from Takara.

### Electroporation of pESCs

1×10^7 cells were electroporated with 20 mg of DNA using a Lonza nucleofector and then seeded on MEFs.

### Karyotyping

Live cells at passage 15 were sent to Cell Line Genetics for cytogenetic analysis. Analysis was performed on twenty G-banded metaphase male pESCs. Two abnormal male clones were analyzed. Clone 1 contained eighteen cells with trisomy 16 and trisomy 17. Clone 2 contains 2 cells which demonstrated trisomy 16.

### i-GONAD

Mouse oviduct electroporation was performed as previously reported (Ohtsuka et al., 2018). Briefly, 8-16 week old *Peromyscus* were mated with stud males and checked daily for sperm using vaginal lavage. If sperm was found, the female was subjected to iGONAD the same day. CRISPR gene editing cocktails were freshly assembled and contained 6 μM Cas9 protein (IDT 1081058, Lot # 0000405530), 30 μM gRNA, and 18 μM ssODNA (produced in-house). This cocktail was delivered into the oviduct of female mice through microcapillary injection. Oviduct electroporation was performed using a NEPA 21 Electro-Kinetic Transfection System with a range of conditions including: Pd A: between 100-135 mA, Pd on: 5 msec, Pd off: 50 msec, three cycles, decay 10%.

### Genotyping

The screening of founder pups was performed by PCR and direct sequencing using DNA obtained from ear slices. Genomic DNA was extracted from ear slices with the GenElute Mammalian Genomic DNA Mini-prep Kit (Sigma-Aldrich). PCR was performed with PrimeSTAR® DNA Polymerase (Takara Bio) and the following primers *Rosa26-F (5’-GGGAGTTCTCTGCTGCCT-3’) and Rosa26-R (5’-AGTTTTGCTGCATAAAACCCC-3’)*.

### Temperature & Activity Data Monitoring

Data were gathered with G2 E-Mitter implants for core body temperature and locomotor activity (Starr Life Sciences Corp., Oakmont, PA). G2 E-Mitters were implanted in the abdominal cavity under anesthesia using a cocktail containing ketamine (150mg/kg), xylazine (5mg/kg) and acepromazine (2mg/kg). E-Mitters were sutured to the ventral muscle wall to maintain consistent core temperature measurements. Recordings began immediately, but mice were not mated until at least 3 cycles of regular estrus were observed. Recordings were collected continuously at 1-min resolution for core body temperature and activity. All mice were between 10 and 16 wk of age at the time of implant surgery and were handled once/wk across recordings at the time of cage changes, but otherwise were left undisturbed in single housing.

### *P. Leucopus* Mating Set-up in Estrous Tracking Experiments

Animals were maintained under 16:8 LD cycle of ∼400 lux (light) to <1 lux red light (darkness), with lights on from 4am to 8pm. Food and water were available *ad libitum*. Mouse temperature and activity profiles were utilized to monitor estrous cycles remotely and mice were paired on the day of apparent estrus from 1 hr before lights-off (7pm) to 5 h after lights-on the following day (9am). Vaginal plugs were not monitored. Removal of the male was the only disturbance to the females.

### *Mus musculus* Estrous Tracking Experiments

8 week old *Mus musculus* (C57BL/6) were maintained under 15:9 LD cycle of ∼400 lux (light) to <1 lux red light (darkness), with lights on from 6am to 9pm. Food and water were available *ad libitum. Mus* activity profiles were used to monitor estrous cycles remotely. Cage changes were the only disturbance to the animals.

### Hamster Estrous Tracking Experiments

14 week old Syrian hamsters *Mesocricetus auratus* were maintained under 12:12 LD cycle of ∼400 lux (light) to <1 lux red light (darkness), with lights on from 7am to 7pm. Food and water were available *ad libitum*. Hamster activity profiles were used to monitor estrous cycles remotely. Cage changes were the only disturbance to the animals.

### Camera Setup, Data Export & Analysis

Each Raspberry Pi Zero 2 W was assigned a unique ID used to organize the data it processed. Each drive had a capacity of 32GB and was flashed with the Raspbian OS image and a script for startup configuration. The Raspberry Pi NoIR Camera V2 modules were connected to the Raspberry Pi Zero 2 W using a ribbon cable, and both were assembled into the official Raspberry Pi Zero case. System monitoring and alerting was done using DataDog and PagerDuty, respectively.

The Raspberry Pi systems were attached to acrylic stands using Command strips for ease of repositioning. The stands were placed next to each cage. For one experiment, near-IR (940nm), 100mA LEDs were attached to the stands and powered by the Raspberry Pi USB port to illuminate the mice at night.

Linux’ systemd utility was used to manage the image capture, processing and upload services. The image capture service captures 640×480 RGB images every 4 seconds using raspistill and saves them locally as jpegs. Every 10 minutes, the image processing service runs a Python program that uses OpenCV to compare each 2-tuple of consecutive images using the imagediff algorithm. Next, this service attempts to upload this data to a Postgres database in AWS RDS, then deletes the processed images to save storage. To mitigate network reliability issues, a copy of the data is written to a local file, to be recovered when the connection is restored. If enabled, the image upload service uploads stills to AWS S3, and inserts a corresponding row referencing the S3 URI into the database. Stills are stored until a successful upload, or until the disk reaches capacity.

All Raspberry Pis are connected to a VPN running on an AWS EC2 instance for remote debugging and to receive over-the-air software updates; other data was not sent over the VPN. Pis uploaded their IP address to S3 every 2 minutes, in part serving as a health check.

An AWS EC2 instance runs an alerting system as well as a web server for monitoring and data visualization. Every 6 hours, a Python script checked that each Pi has uploaded its IP and data recently, and sent an email listing the Pis that have not. Every hour, the Raspberry Pi ID to mouse ID mapping was read from a Google Sheet and inserted into the database. Data was then downloaded from the database per mouse, grouped into 5-minute windows, and stored in a day by window format. Further data processing in the form of outlier removal was available interactively on the web interface.

All will be made available upon publication, or request.

### Vasectomies

Briefly, 8-12 week old male *Peromyscus* were anesthetized with a cocktail containing ketamine (150mg/kg), xylazine (5mg/kg) and acepromazine (2mg/kg) in preparation for surgery. Once anesthetized, the eyes were covered with ointment to prevent dryness and the abdomen was shaved with small clippers and cleaned with betadine followed by 70% ethanol, 3 times. Following aseptic technique, a small incision of about 1.5cm was made at a point level with the top of the legs. Another small incision was made in the body wall along the linea alba. With blunt forceps, the fat pad was pulled out, exposing the vas deferens. The vas deferens was cauterized with a high temperature cautery power handle. The fat pad was then reinserted inside the body wall, and repeated on the other side. The body wall was sutured with 1-2 stitches of coated VICRYL sutures and the skin was closed with wound clips which were removed 7-14 days later.

### Embryo Transfers

Briefly, 8-16 week old female Peromyscus were mated with vasectomized males on the night of apparent estrus to generate pseudopregnant recipients. The following day, females were anesthetized with a cocktail containing ketamine (150mg/kg), xylazine (5mg/kg) and acepromazine (2mg/kg) in preparation for surgery. Once anesthetized, the eyes were covered with eye ointment to prevent dryness and the lower back of the recipient mouse was shaved with small clippers. Using cotton tip applicators, the surgical area was cleaned with betadine followed by 70% ethanol, repeated 3 times. Following aseptic technique, a small transverse incision (less than 1cm) was made with fine dissecting scissors about 1cm to the right of the spinal cord at the level of the last rib. Once the ovary or fat pad could be visualized through the body wall, a small incision was made just over the ovary. With blunt fine forceps, the fat pad was pulled through the incision and the ovary, oviduct and uterine horn was pulled out and laid down over the middle of the back, so that the oviduct was exposed outside the body wall. Manipulated 2-cell embryos were transferred into the oviduct by puncturing the oviduct wall, in between the infundibulum and ampulla with a sterile 30g needle and transferring the embryos through the small opening. Once the embryos have been transferred, the ovary, oviduct and uterus were placed inside the body wall. The body wall was sutured with 1 or 2 stitches of coated VICRYL sutures and the skin was closed with wound clips, which were removed 7-14 days after embryo transfer.

### Pronuclear Injection

Briefly, Peromyscus females were used as embryo donors and recipients. Camera-tracked donors were mated with proven studs. Two days later, 2-cell embryos were flushed from the oviducts using prewarmed M2 medium. After embryo collection, 2-cell embryos were transferred to pre-equilibrated KSOM (CytoSpring; K0102) overlaid with mineral oil and incubated at 37° C (5% CO2 and 5% O2). SpCas9 protein (final concentration 50 ng/ul) and sgRNA (final concentration 50 ng/ul) were mixed and incubated at room temperature for 15-20 minutes. DNA template was added to the RNP solution at a final concentration of 10 ng/ul. The pronuclei of each blastomere was injected for each 2-cell before incubating the embryos in KSOM. Pressure for microinjection was provided by Eppendorf FemtoJet 4i. Peromyscus females were mated to vasectomized Peromyscus males one day prior to 2-cell microinjection. The same day as microinjection, manipulated 2-cell embryos were transferred to pseudopregnant females’ oviducts.

### Image Processing For *Mus* and Hamsters

Still images captured every 4 seconds with a resolution of 640×480 were cropped to retain only the quadrant where food intake activities were observed. For *Mus*, only images from 0600-0900 UTC each day were used for downstream analysis. For hamsters, only images from 0000-0820 UTC were used. Each frame was then normalized to have a minimum of 0 and maximum of 255 in each of the 3 channels. “TwoPoint” background subtraction algorithm from Python package bgslibrary was then applied iteratively to each frame and a background learned and subtracted from each frame. These frames were then normalized to a range of 0 to 1, and L2 norms computed for each frame and stored in a separate array for each day. To reduce the impact of spurious activities, adjacent elements in this array above a threshold of T but forming blocks within the index range of (p,q) of less than N or more than N’ neighbors were replaced with the mean of elements with indices p-1 and q+1. For *Mus*, T=25, N=20, N’=200. For hamsters, T=15, N=3, N’=100. The differences between adjacent elements of this resulting array was then computed for each day and plotted as a heatmap. For heatmap visualization, a max and min normalization value was set using 30 break histogram binning. More specifically, a 30 bin histogram of the log10 difference was generated with 0 and 1 set just outside the tails. For the cumulative activity plots, activity levels for each time point were convolved with a kernel of one hundred ones, max pooled with a block size of 100 and the daily sum was plotted as time series bar plots. Autocorrelation of daily activity levels was then computed to identify periodicity with a 95% confidence interval.

### Thermal Activity Tracking

Still images were captured using the FLIR Lepton 3.0 (Teledyne FLIR, Wilsonville, OR) attached directly to a Raspberry Pi Zero 2 W computer or to a PC via a PureThermal 3 adapter (GroupGets, Reno, NV). An IR-transparent zinc selenide disk was used to construct an imaging window for the thermal camera. Images were sectioned according to the Image Processing For Mus and Hamsters section above. Cropped images were then binarized and the difference between the L2 norms of two adjacent frames was used as a proxy of activity for downstream analysis. Again for heatmap visualization, a max and min normalization value was set using 30 break histogram binning. For the cumulative activity plots, activity levels for each time point were convolved with a kernel of one hundred ones, max pooled with a block size of 100 and the daily sum was plotted as time series bar plots. Autocorrelation of daily activity levels was then computed to identify periodicity with a 95% confidence interval.

### Homology Arm Sequences for Targeting Constructs

#### 5’ Rosa26 Homology Arm

GGTGGTCAGGGCCATGGTCCGCCCTGCCGCAACCGGAGGGGGTGGGAGGAGGGAGCGGAAAGATCTCCACCGGACGCGGCCATGGCTCCCACGGGGGGCGGAGGAGCGCTTCCGGTTCCCGTCTCGTCGCTGATTGGCCTTTTCTCCTCCCGCCGTGTGTAAAAACACAAATGGCGTGTTTTGGTTGGAGTGAAGCGCCTGTCAGTTAACGGGTGCTGGAGTGCGCAGCCGCTGGCTGCCTCGCTGTGCCCACTGGGTGGGGCGGGAGGTAGGTGGGGTGAGGAGAGCTGGACGTGCGGGCGCGGCCGGCCCCTGGCGGGGCGGGGGAAGGGAGGGGAGGGAGGGTCAGCGAAAGTGGCTGGCGCCCAGGCGGCCTCCCACCCTCCCCTTCCTCTGGGGGAGTCGGTTTACCCGCCGCCGGCCTGGCCTCGTCGTCTGATTGGCTCCCGGGGCCCAGAAAACTGGCCCCTGCAATTGGCCCGCGTTCGTGCAAGTTCAGTCCATACGCGGGCTGGCGGGGGCGGCAGGGAGGCGCTCGCAGGTTCCGGCCCTCCCCAAGGCCCCGCGCCGCAGAGTCTGGCCCCGCGCCCCTGCGCAACGTGGCAGGAAGCGCGCGCTGGGGGCGGGGACGGGCAGGCGGTCTGAGCGGCGGGGCGGGGGCGAGCGCGGTTCCTCCTAGAGTTGTTGCCGCACGAGGGGCGAGCTGAACCGGACCTGCATGGCGCACTCCTAGGAGTGGAGGAAGGAGTGGGGGCTCAGTCGGGCCGGTTTGGAGGCAGGAGGCACTTGCTCTCCAAAATTCCGTGAGAGCTGCAATCAATAAGGGAGCCGCAGTGGAGTAGGCGGGGAGAAGGCCGCACCCTTCTCGGCAGGGGGAGGGGAGTGCCGCAATACCTTTCTGGGAGTTCTCTGCTGCCTCCTGGCTTCTAAAGACCGCCCCGGGCCTGGAACAGTCCCTTCCCCCTCTCCCCTCGTGATCTGCAAGTCGAGGCTTTCTGGA

#### 3’ Rosa26 Homology Arm

AGATGGGCGGGAGTCTTCTAGGCTGGCTTAAGGGCTAACCTGGTGCGTGGGCGTTGTCCTGCAGGGGAATTGAACAGGTTTAAAATTGGAGAAACGACACTTCCCACAGATTTTCGTTTGTGTCCGGAGGGAATTGTAATAGGAGCAAAGGAGGGAAATGGGTGACTAGGCGCTCACCTGGGGTTTTATGCAGCAAAACTACAGGTTATTATTGCTTATGATCTGCCTTGAAGTATTTTTCATCGTGTTAGATTAAAGACATGCTCACCCAAGTTTATACTCCCACTTGAGATCCTTGTTATATCATGAAATTTTAGTATCGTAAATTAGAATATATAAACAGAATCTTAGCAGTTTCTGCAGAGCCCAGTGCTTCACTCTGTCCCTCTTGCTCCCTCTGCAGCCCTACCAAGAGATATTTTAGCGCTCTCTCATTTTAGTCCCGTTTTCATTTGTTGTACTGGCTTATCCAATCCCTAGACAGAGCACTGGCATTCTTCCTCTCCTGATCTTAGAAGCCTGATGAGTCATGAAACCAGACAGATTAGTTACACCACAAATTGAGGCTGTAGCTAGGGCCTTGCCCTGCAGTTCTTTTATTCCTCCTTAGTACATTTTGTTGACTGTTTGCCTTGATTTCATTTTCTATCCCCACCCTTCAGGAGCTTTACTACAATAGATTTATTGAGAATCTAACCTAAAGTTCAGCTTCTTTCCACTCCCATGTATGCCTATCTCCTTTTTCACCATTAATAAACTGAACTGTTATTAATGGCTTCCAGATGGATGTCTCCTCCAATATCACCTGATGTATCTTACATATCGTCAGGCTGATTGTATTAAAAGGTACATGAATTGACATGAAATTTACTGACTCCTTAATGCTTTAGTGGATTCCATGAGTACAATACAGAAGACCGTTAATGGGCTACTGACAGCTTTTTCATTTCATTATTTCCTGCATACACACTTTGAGTCACA TGGAATTGATTACTGCTTA

