## Supplement for "Low-cost camera-based estrous tracking enables transgenesis in *Peromyscus leucopus*, the primary reservoir for Lyme disease"

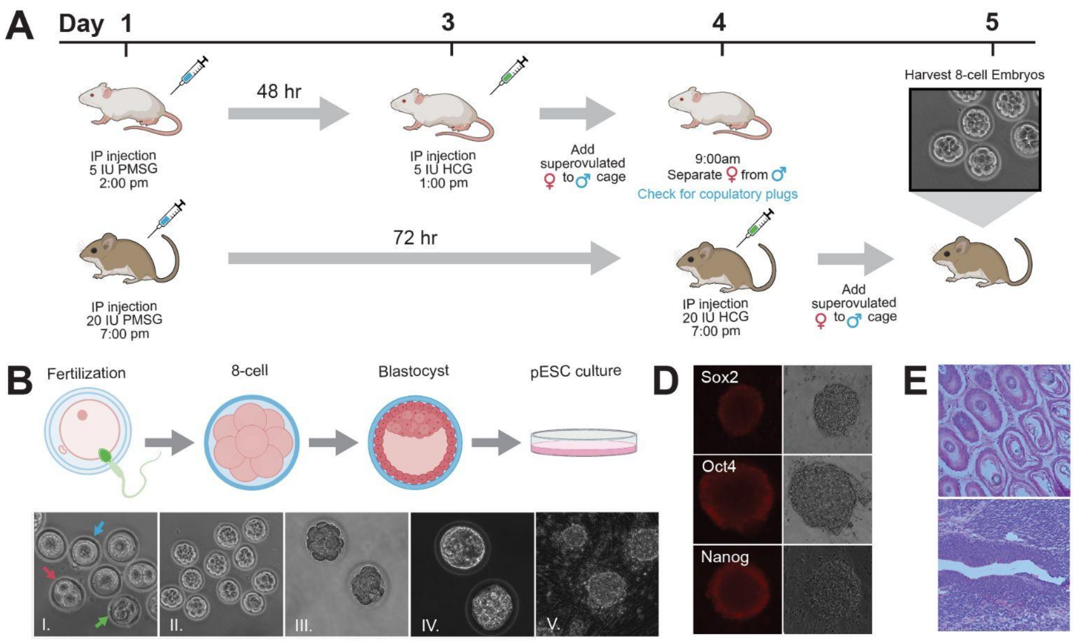


C

| Attempt | Blastocysts (# Isolated) | Number of Outgrowths | # ESC colonies | Media | Outcome | Citation |
|---|---|---|---|---|---|---|
| 1 | 2 | 1 | 0 | 15% KSR PD98059 + Mouse LIF | ESC-like colonies present for < 5 passages. Slow growth and no expansion. | Markoulaki et al, 2008. Methods |
| 2 | 4 | 1 | 0 | 15% KSR PD98059 + Mouse LIF + 2i | Passaged < 3 times. No stable lines achieved. | Hanna, Markoulaki et al., Cell Stem Cell, 2009. |
| 3 | 4 | 0 | 0 | 15% KSR + PD98059 + Mouse LIF + 2i + N2B27 | No pESC colonies formed after dissociation*. | Buehr et al, 2008, Cell |
| 4 | 7 | 2 | 0 | 15% FBS PD98059 + mouse LIF + 2i | No pESC colonies formed after dissociation*. | |
| 5 | 9 | 9 | 5 | 15% KSR PD98059 + Mouse LIF + 2i | Clones are challenging to maintain long-term as pESCs and often differentiate. | Hanna, Markoulaki et al., Cell Stem Cell, 2009. |

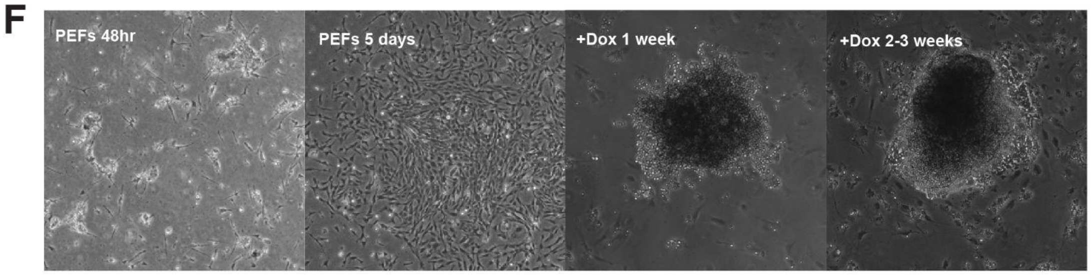


**Supplemental Figure 1. *Peromyscus* ESC and iPSC Derivation**

**A)** Diagram of *Mus* and *Peromyscus* hormone priming protocols. **B)** *Peromyscus* embryo development and ESC derivation. **C)** Table of ESC culture conditions and outcomes. **D)** ESC colonies stained positive for pluripotency markers: Sox2, Oct4, Nanog. **E)** Cryosections from teratoma formation assay. Top: mesoderm layer. Bottom: ectoderm layer. **F)** iPSC attempts. 1st image: Embryonic fibroblasts 48 hours *in vitro*. 2nd image: Embryonic fibroblasts 5 days *in vitro*. 3rd image: 1 week after adding doxycycline to induce expression of reprogramming factors. 4th image: 2-3 weeks after adding doxycycline to induce expression of reprogramming factors.

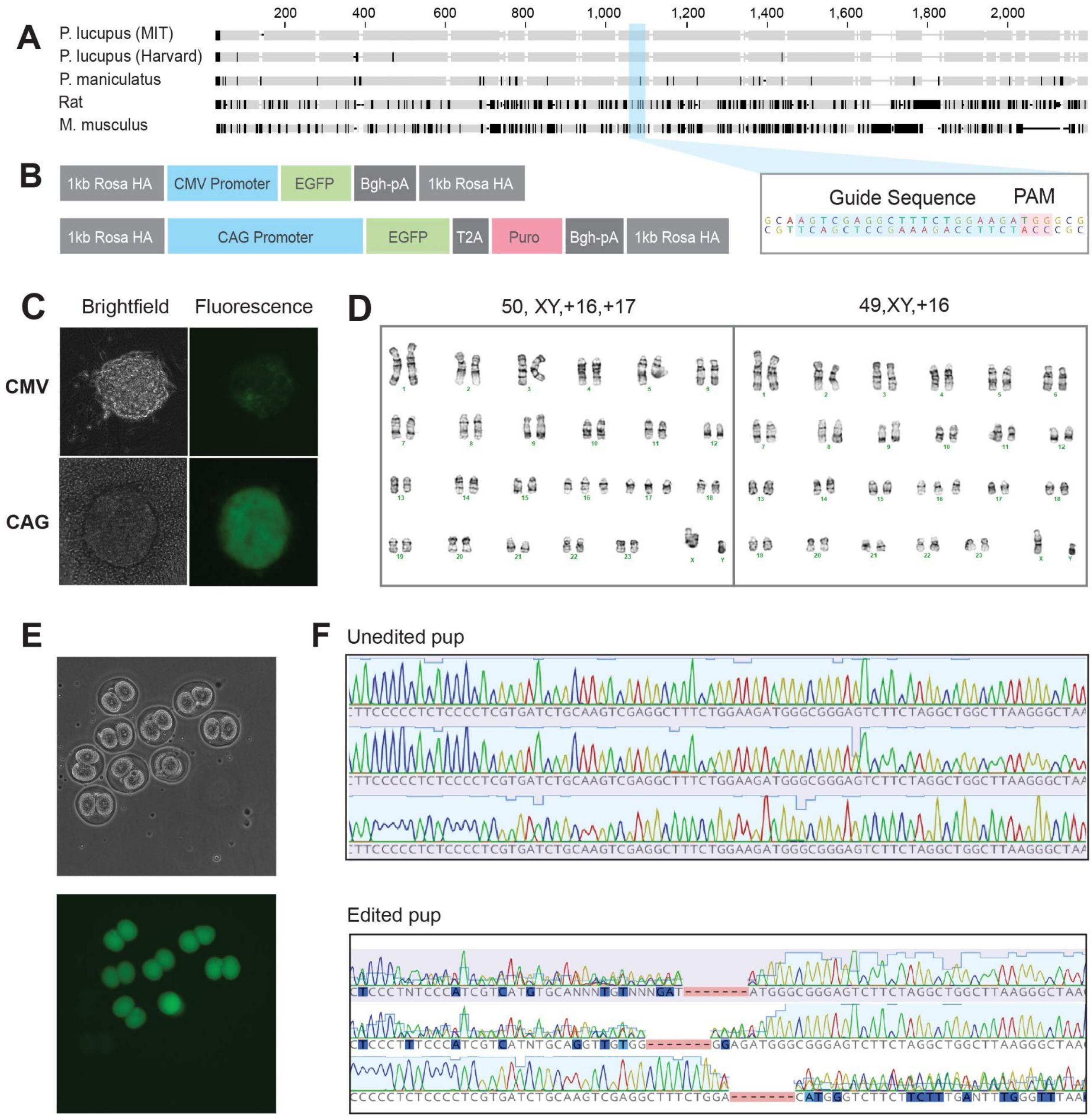


**Supplemental Figure 2. Engineering *Peromyscus* Embryos**

**A)** Alignment of Rosa26 locus from *de novo Peromyscus leucopus* genome with multiple genomes from the UCSC Genome Browser. The *Peromyscus leucopus* Rosa26 guide sequence is highlighted. **B)** *Peromyscus leucopus* Rosa26 targeting construct designs. **C)** pESC colonies. Top: Colony targeted with the pR26-CMV-EGFP construct. Bottom: Colony targeted with pR26-CAGGS-EGFP-2A-Puro construct. **D)** Abnormal karyotypes from two pESC colonies. **E)** Detection of eGFP fluorescence in 2-cell *Peromyscus* embryos following delivery of eGFP mRNA via i-GONAD. **F)** Sequencing reads from wild type and mutant *Peromyscus* pups from mother subjected to i-GONAD. The mutant pup has a mutation in the Rosa26 locus.

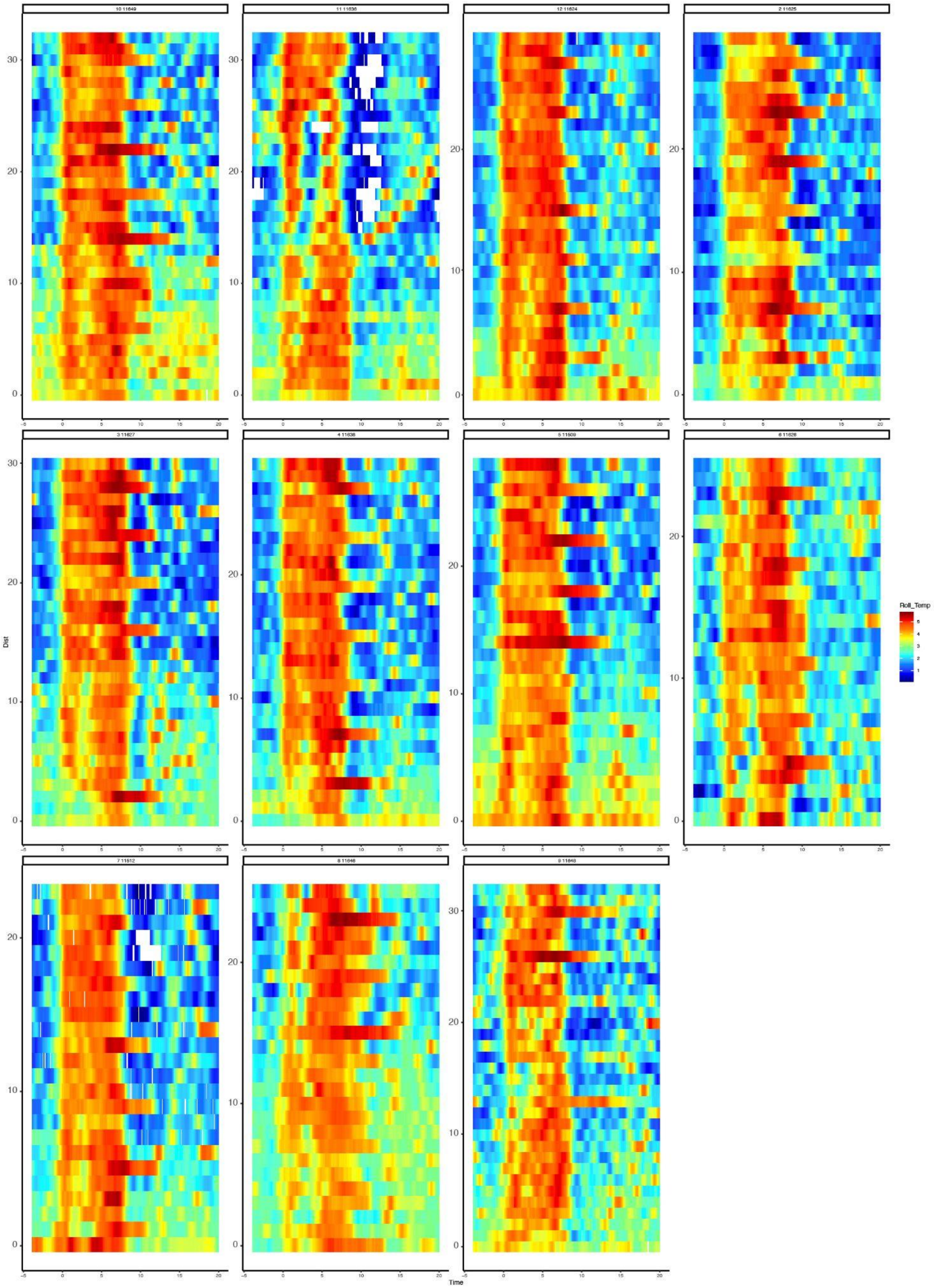


**[Supplemental Figure 3](). Estrous Cycling Following Surgery**

Temperature graphs for *Peromyscus* with implanted sensors. Cycling patterns could be observed 3 to 14 days after sensor implantation surgery. X axis: 0 = 8pm when lights turn off in the animal facility. Y axis: 0 = first day after surgery.

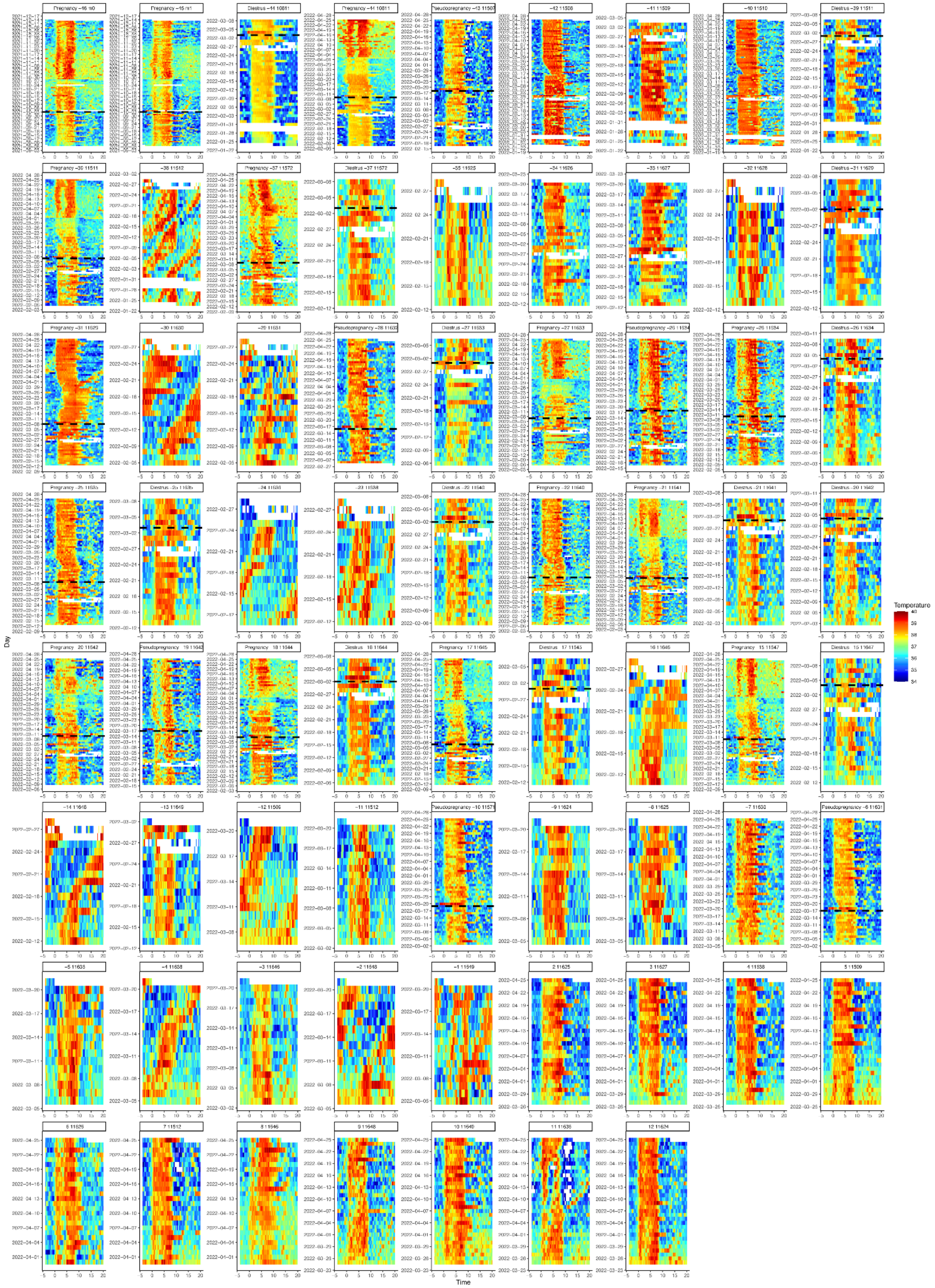

**Supplemental Figure 4. Temperature Graphs for Sensor Mice**

Temperature graphs for all *Peromyscus* with implanted sensors, including mice from ongoing experiments. X axis: 0 = 8pm when lights turn off in the animal facility. Y axis: sequential days stacked from bottom to top.

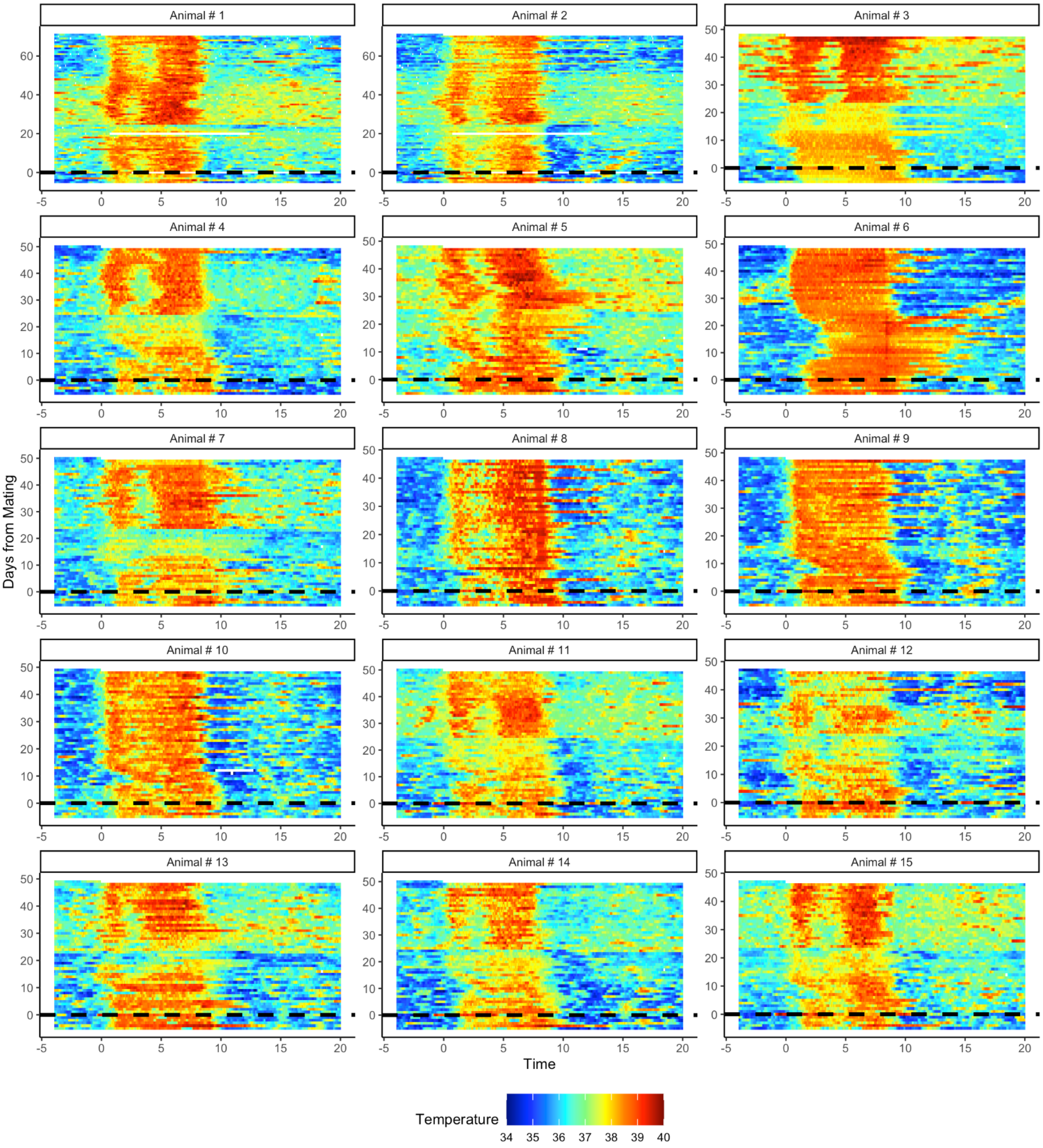


**Supplemental Figure 5. Temperature Tracking of 15 Mice Mated in Estrus**

Temperature graphs for *Peromyscus* before and after timed mating. Mice with implanted sensors were tracked to determine the timing of their estrous cycles and then mated with proven studs on the night of presumed estrus. X axis: 0 = 8pm when lights turn off in the animal facility. Y axis: 0 = day female *Peromyscus* were mated with stud males.

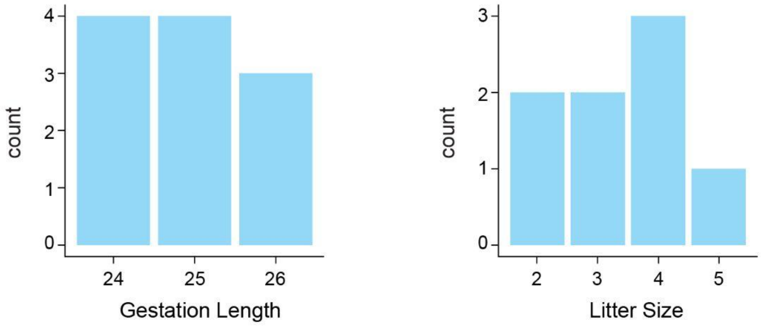


### [Supplemental Figure 6](#). Timed Mating Results

Gestational length and litter size for 11 mice that produced pups after timed mating. While the date of birth was noted for all 11 litters, pups were only counted in 8 of the 11 litters, as litters were cannibalized.

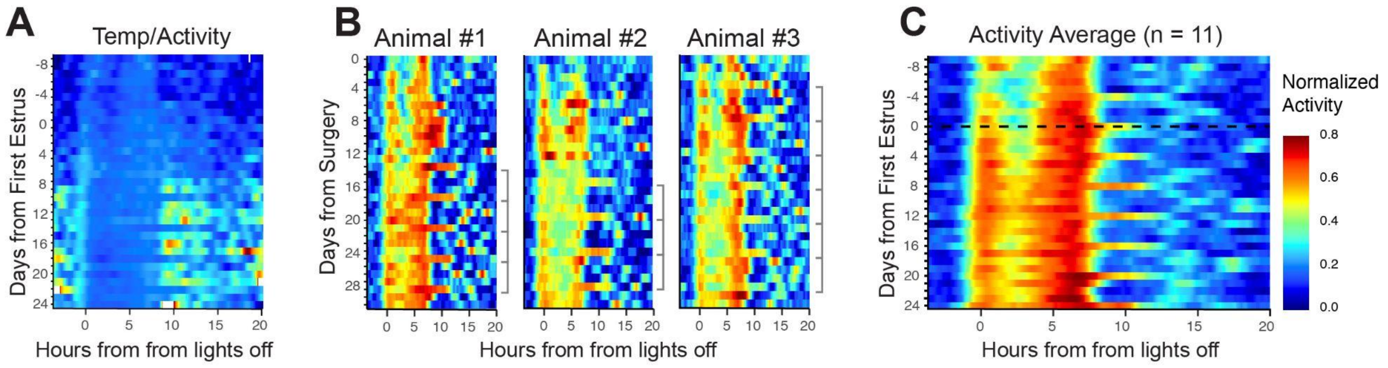


### [Supplemental Figure 7](#). Sensor Activity Data

**A)** The ratio of core body temperature and activity shows that temperature readouts are highly correlated with corresponding activity data. **B)** Activity data from three individual female *Peromyscus*. Prolonged extensions of activity (estrus) coincide with elevated temperatures. X axis: 0 = 8pm when lights turn off in the animal facility. Y axis: 0 = first day after surgery. **C)** Mean activity for 11 female *Peromyscus* shows activity consistently aligns with temperature trends. X axis: 0 = 8pm when lights turn off in the animal facility. Y axis: 0 = first estrus day after surgery.

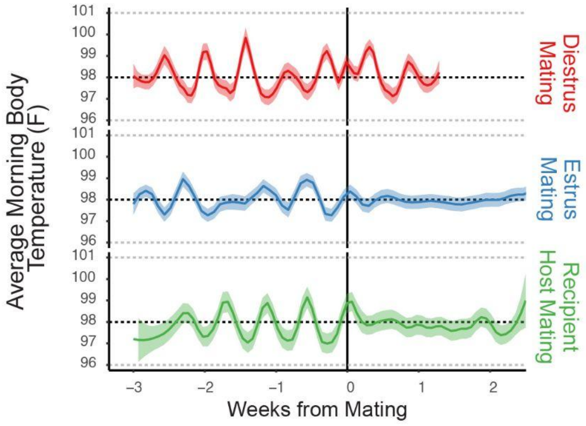


### [Supplemental Figure 8](#). Comparison of Mating Outcomes

Morning core body temperature for pregnant and pseudopregnant *Peromyscus* is similar, though the suspension of the estrous cycle is truncated in pseudopregnant mice. No disruption of morning temperature cycling is observed when *Peromyscus* are mated in diestrus.

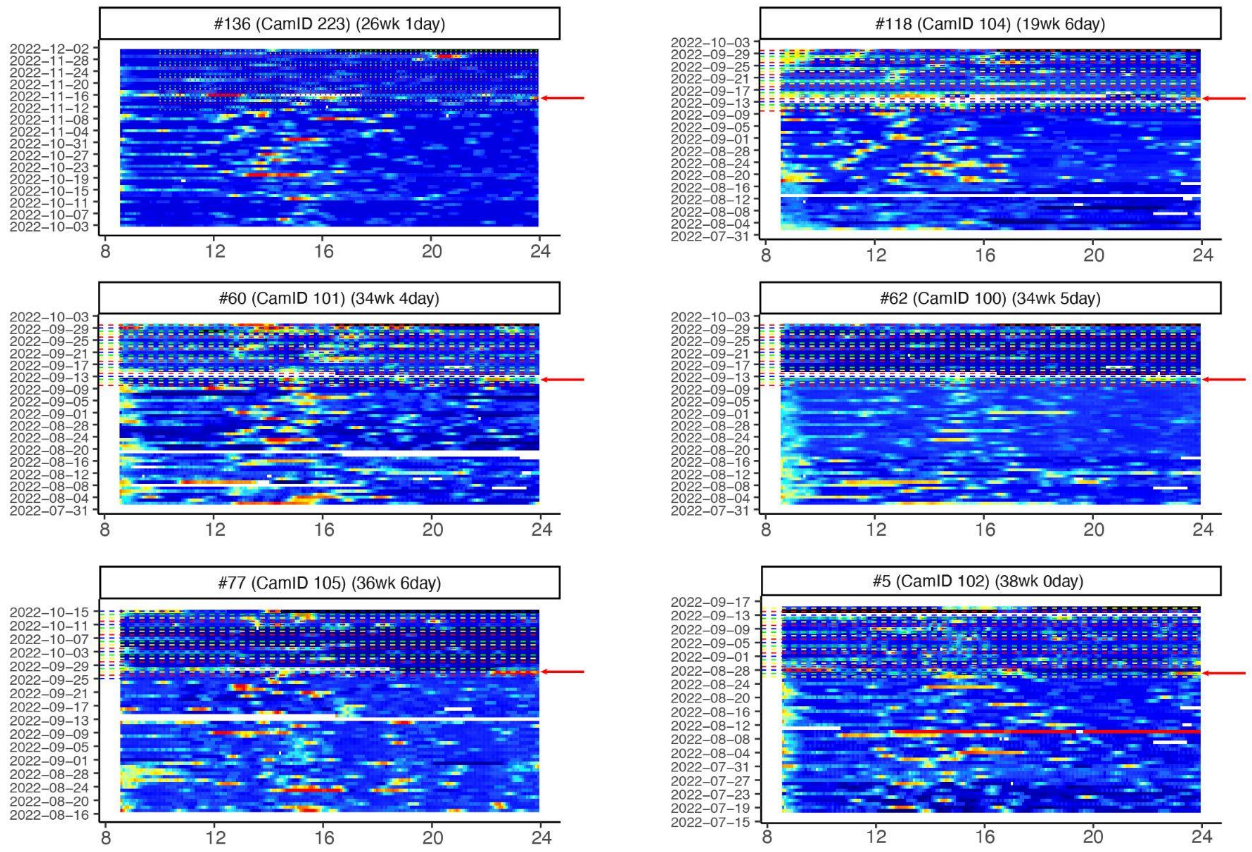


**[Supplemental Figure 9](#). Shifted Estrus Day after Timed Matings with Vasectomized Males**

Activity graphs from camera-tracked mice before and after timed mating with vasectomized males. Red arrows point to the evening females were mated. All 6 mice continued to cycle post mating with cycles shifted by one day. X axis: 0 = 8pm when lights turn off in the animal facility. Y axis: sequential days stacked from bottom to top.

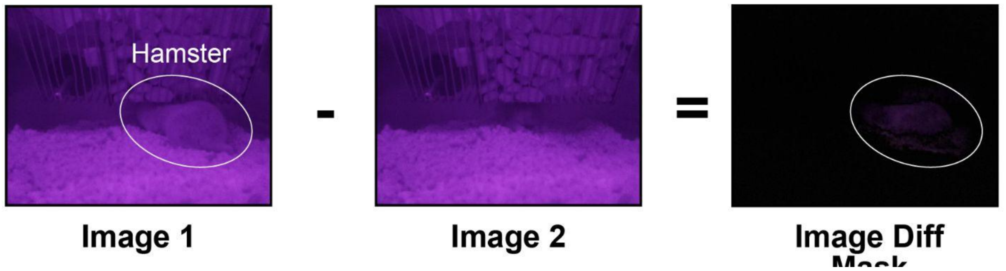


**[Supplemental Figure 10](#). Image Difference Analysis**

The original camera-based activity tracking system applies pixel-wise subtraction on two successive images and then sums the absolute values of the differences to produce an activity metric. However, if two successive images were similar in color when under IR light (like image 1 and image 2), the resulting image difference mask can fail to capture animal activity.

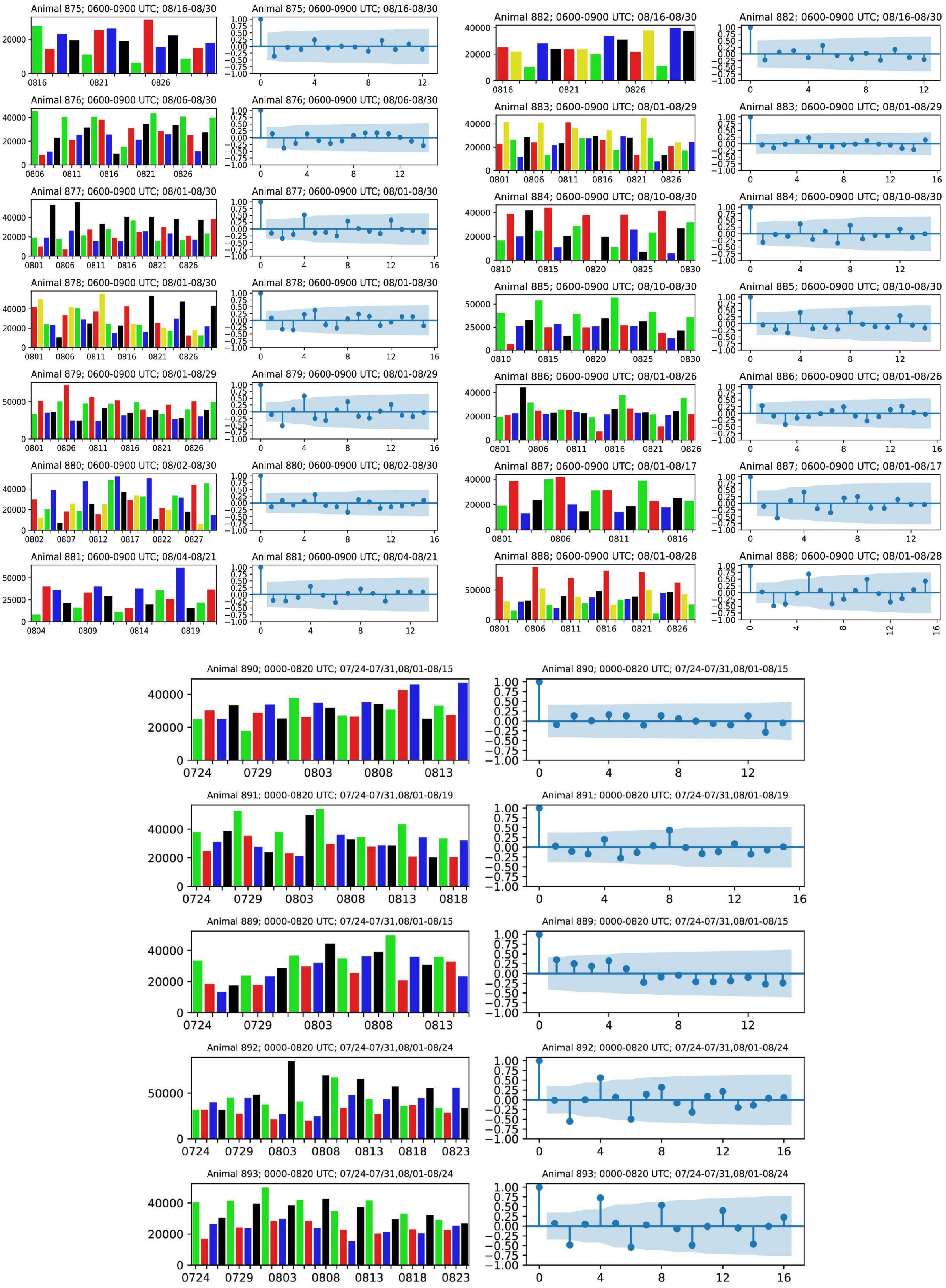


**Supplemental Figure 11**. ***Mus* and Hamster Cumulative Activity Plots & Autocorrelation**

Top left and right: Cumulative activity for 14 *Mus* with either 4 or 5-day color schemes, and corresponding autocorrelation post sectioning and background removal; autocorrelation shown with a 95% confidence interval. Bottom: Cumulative activity for 5 hamsters with 4-day color scheme, and corresponding autocorrelation post sectioning and background removal; autocorrelation shown with a 95% confidence interval.

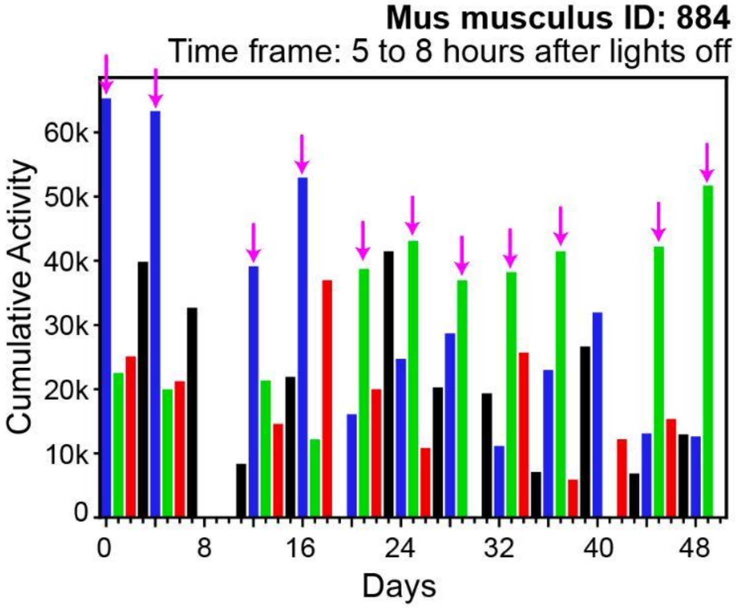


**Supplemental Figure 12**. ***Mus musculus*** **with Shifting High Activity Day**

Daily cumulative activity plot for *Mus musculus* with shifting high activity day. Purple arrows point to high activity days, which appear to shift from a 4-day "blue" cycle to a 4-day "green" cycle. X axis: daily cumulative activity 5 to 8 hours after lights off. Y axis: sequential days from left to right.
